# Benchmarking Generalizability in Deep Learning-Based White Matter Tract Segmentation

**DOI:** 10.64898/2026.07.30.741597

**Authors:** Junbeom Kwon, Gabriele Amorosino, Franco Pestilli

## Abstract

White matter tracts (WMTs) are the brain’s structural foundation for information transfer, underlying essential cognitive and behavioral functions. While diffusion MRI and tractography enable non-invasive mapping of these pathways, automated segmentation often lacks generalizability across diverse data sources. We conducted a systematic, cross-dataset evaluation of four state-of-the-art deep learning architectures, benchmarking their performance across independent datasets with varying acquisition protocols and populations. CNN-based models such as TractSeg achieved the highest within-domain accuracy, but performance dropped sharply under domain shift, most severely when we applied adult-trained models to pediatric data. To address this degradation, we introduce Ensemble White Matter Tract Segmentation (EWMTS), which combines complementary models to partially recover accuracy under domain shift, although performance still falls short of within-domain levels. By openly releasing this benchmark and a reproducible processing pipeline, we provide the neuroimaging community with a framework to develop and benchmark segmentation models across the heterogeneity of real-world neuroimaging data.

## INTRODUCTION

Myelinated axon bundles form the brain’s white matter and enable fast, reliable signal transmission across distributed neural circuits. These organized pathways, collectively termed white matter tracts (WMT), serve as the brain’s information highways, underlying all major behavioral, cognitive, and perceptual processes ^1^. WMT properties mature during development and degrade with aging, with developmental rates predicting later degeneration ^2^. Disruptions in WMT characterize numerous disorders, from autism spectrum disorder ^3,4^ to cerebral small vessel disease ^5^. Therefore, accurate WMT characterization is essential for elucidating brain network architecture, investigating structure-function relationships across populations, and improving clinical assessment of neurological and psychiatric disorders ^6–8^.

Diffusion-weighted magnetic resonance imaging (DWI) combined with computational tractography ^8–11^ enables non-invasive reconstruction of WMT. The Segmentation of WMT (hereafter referred to as WMTS) is a complex process that requires integrating anatomical knowledge into a computational model. Traditionally, WMTS relied on manual delineation of anatomical regions of interest (ROIs) by experts to filter tractography streamlines ^12^, but the substantial time and expertise required have motivated the development of automated methods. Traditional automated WMTS methods fall into two categories: atlas-based and clustering-based approaches. Atlas-based methods register individual data to templates and delineate tracts using predetermined ROIs ^13,14^ or expert-labeled fiber templates ^15^. Clustering-based methods group streamlines using unsupervised algorithms based on proximity and similarity ^16,17^. However, atlas-based approaches suffer from registration inaccuracies and false positives, while clustering-based methods struggle with inconsistent anatomical classification ^18^.

Recent approaches use deep learning (DL) methods to perform WMTS. Two DL-based WMTS approaches have been proposed: streamline-based and voxel-based. Streamline-based approaches segment WMTs by using streamline features and matching streamlines to each WMT. Streamline-based approaches evolved from early CNN approaches ^19^ to more sophisticated architectures, including Graph Convolutional Networks ^20^, point-cloud frameworks ^21–25^, and Transformers ^26^ that capture complex fiber geometry. Voxel-based approaches instead segment the brain volume that overlaps with each WMT ^27^. Currently, voxel-based segmentation dominates clinical applications due to superior processing speed, lower computational complexity, and costs. TractSeg was the first model to use volume-based approaches ^28,29^. **Fig. 1** illustrates this voxel-based pipeline, in which fiber orientation distribution function (fODF) peak maps serve as input to a segmentation model that outputs a binary volume mask for each WMT. Since TractSeg, a variety of other models have been proposed, all of which use manually annotated data for training. These models focus on DWI data ^30–34^, or multimodal inputs ^33,35–37^. While these models predominantly rely on manually annotated data, self-supervised learning has been explored to reduce annotation dependency ^38,39^, and transfer learning strategies have been employed to generalize models across novel WMTs ^35,40–43;^ see also ^44–46^ for reviews).

**Figure 1.**
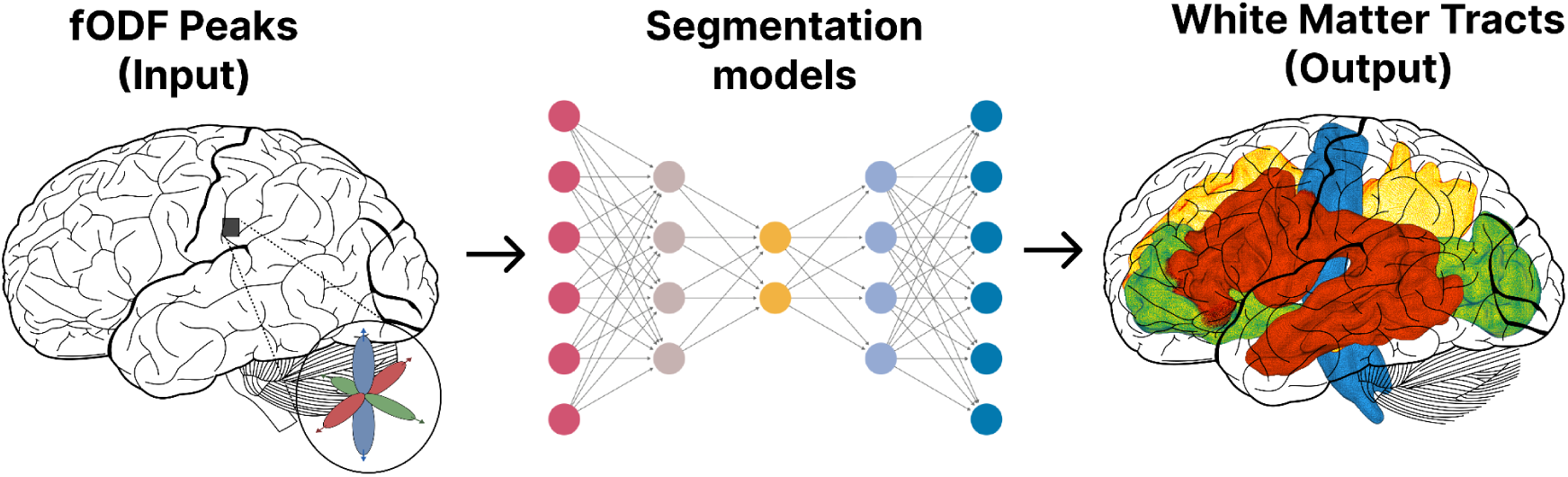
Illustration of the voxel-based white matter tract segmentation. Fiber orientation distribution function (fODF) peak maps derived from diffusion MRI (left) serve as input to a deep learning segmentation model (middle). The model outputs volumetric white matter tract (WMT) predictions (right), producing a whole-brain segmentation in which each color represents an anatomically distinct major WMT, including the Arcuate Fasciculus (red), Cingulum (yellow), Corticospinal Tract (blue), and Inferior Fronto-Occipital Fasciculus (green).

Despite substantial advances in deep learning (DL)-based white matter tract segmentation (WMTS), the development of robust and generalizable methods remains constrained by three interrelated challenges spanning data, evaluation, and model behavior. First, limitations in training data, specifically insufficient dataset size and variability, restrict the ability of DL models to learn representations that capture the full spectrum of anatomical diversity, acquisition protocols, and population differences; large-scale, representative datasets remain scarce ^44,47^. Second, benchmark and evaluation frameworks are underdeveloped: despite the proliferation of proposed models, only a single comprehensive benchmark has systematically compared DL-based WMTS approaches ^32^, leaving performance across heterogeneous datasets largely unquantified. Third, as a consequence of these limitations, current models exhibit poor generalization under domain shift, with performance degrading when applied to data that differ from the training distribution due to variations in scanners, preprocessing pipelines, or anatomy ^43,48,49^. Although strategies such as self-supervised learning and transfer learning show promise for mitigating these issues, substantial cross-dataset performance gaps persist ^40,43,50,51^.

We address these gaps and make three contributions. First, we share a larger benchmarking dataset spanning 3 scanner types, scanning sequences, and age ranges. Furthermore, we share code and web services for a reproducible preprocessing pipeline that generates our benchmark datasets using *brainlife.io* ^52^. Second, we use a comprehensive, cross-dataset evaluation of state-of-the-art WMTS models, benchmarking modern architectures including hybrid CNN-Transformer models (MedNeXt ^53^, SwinUNETR ^54^, and a foundation model for medical image segmentation (MA-SAM ^55^). Our large-scale unified pipeline enables evaluation of model scalability previously difficult with limited samples. Finally, we propose Ensemble White Matter Tract Segmentation (EWMTS), which combines predictions from multiple models to achieve optimal performance. Together, these advances provide the neuroimaging community with robust, generalizable tools for studying brain connectivity across diverse populations, enabling more reliable investigations of structure-function relationships in health and disease.

## RESULTS

### Benchmark dataset generation, curation, and sharing

Training deep learning models for neuroimaging requires consistent labels paired with data that capture realistic variability in scanner hardware, acquisition protocols, and populations. Without such variability, models may overfit the data, capture primarily dataset-specific structure and show limited transfer performance ^56^. When robust labels accompany realistic data variability, models learn more generalizable representations. Current white matter tract segmentation benchmarks, however, rely almost exclusively on a single manually curated dataset of 105 Human Connectome Project subjects (HCP_105_) ^9,28,57^. This small, homogeneous resource has been repeatedly reused despite being generated through time-consuming subject-by-subject manual curation ^30,32,33,38,42,58^. Although HCP_105_ provides near ground-truth quality, its limited scale and uniformity constrain generalization to real-world settings, where data are often lower in quality and cannot be manually curated; accordingly, models trained on HCP_105_ show reduced performance across scanner configurations, particularly in clinical contexts ^44,58^.

To address these limitations, we constructed and released a substantially expanded benchmark of 1,455 subjects by integrating three public datasets: Human Connectome Project ^9,57^(HCP_1057_; 1,057 young adults aged 22 to 35), Cambridge Centre for Ageing and Neuroscience ^59^ (Cam-CAN_288_; 288 adults aged 18 to 87), and Pediatric Imaging, Neurocognition, and Genetics ^60^) (PING_108_; 108 participants aged 3 to 21). Distinct from prior work, tract labels were generated using standardized, semi-automated and reproducible procedures ^52^ (**Table 3**) rather than manual curation, ensuring consistent labeling while preserving variability across preprocessing outputs, scanners, and protocols. In brief, after reconstructing whole-brain tractography, we automatically segmented 61 white matter tracts ^1,52^ using a modified White Matter Query Language ^61^ that combined cortical regions from the FreeSurfer Destrieux 2009 parcellation ^62^ with tract midpoint estimates as inclusion and exclusion criteria, and we then removed spurious streamlines through outlier-rejection procedures ^13^. We release 1.48 TB of derivatives (**Table 1**), shared in Brain Imaging Data Structure format ^63^ and as *brainlife.io* datatypes optimized for model training within dedicated computational infrastructure. The HCP_1057_ and Cam-CAN_288_ derivatives are openly available as a *brainlife.io* Publication, where access is granted after completing the data use agreement directly on the platform, and all derivatives can be analyzed with the apps deployed on *brainlife.io*. Because the PING data cannot be redistributed directly, we instead forward users to the NIMH Data Archive, where the imaging data can be downloaded after registration and acceptance of the terms of use; the corresponding derivatives can then be reproduced with the apps and methods listed in **Table 3**. For all PING subjects, we additionally distribute the affine matrix for aligning them with other datasets.

**Table 1.**
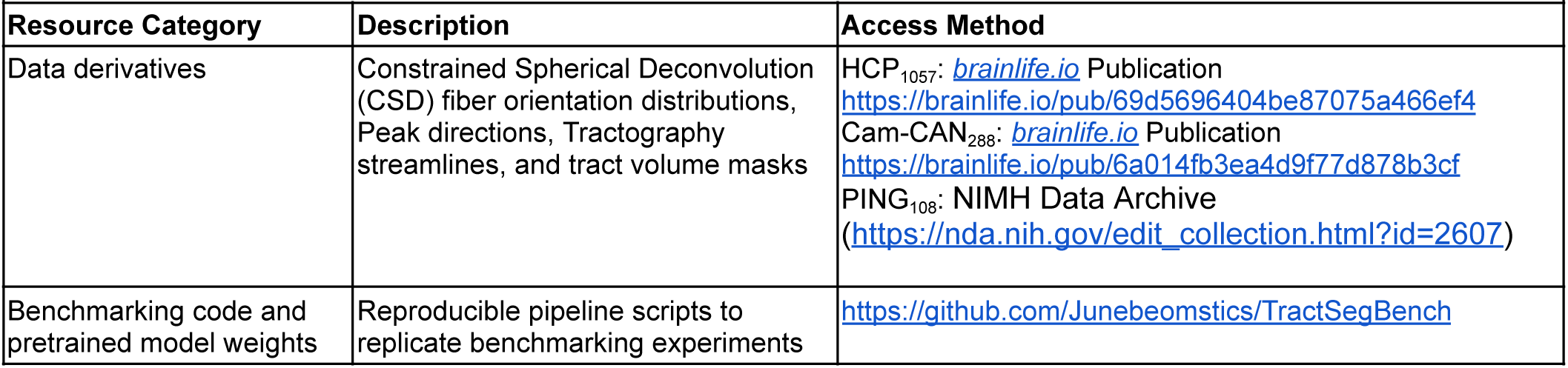
Summary of publicly shared data derivatives, code repositories, and processing applications.

**Table 2.** Datasets used for the study.

| Table 2. Datasets used for the study. |  |  |  |
| --- | --- | --- | --- |
|  | HCP S1200 | Cam-CAN | PING |
| Participants | 1,057 | 288 | 108 |
| Age Range (years) | 22–35 | 18–87 | 3–21 |
| Age (mean ± SD) | 28.7 ± 3.7 | 54.3 ± 18.6 | 12.1 ± 5.2 |
| **Anatomical MRI** |  |  |  |
| Scanner | 3T Siemens Connectome | 3T Siemens Tim Trio | 3T Siemens/GE/Philips |
| Resolution (mm<sup>3</sup>) | 0.7 isotropic | 1.0 isotropic | 1.2 × 1.0 × 1.0 |
| Sequence | MPRAGE T1w | MPRAGE T1w | MPRAGE T1w |
| **Diffusion MRI** |  |  |  |
| Resolution (mm<sup>3</sup>) | 1.25 isotropic | 2.0 isotropic | 2.0 isotropic |
| b-values (s/mm<sup>2</sup>) | 1000, 2000, 3000 | 1000, 2000 | 1000 |
| Directions per shell | 90, 90, 90 | 30, 30 | 30 |
| TE/TR (ms) | 89.5/5520 | 106/9100 | 88/8000 |

**Table 3.** brainlife.io applications (Apps) used in the study. Comprehensive list of the brainlife.io Apps, their associated Digital Object Identifiers (DOIs), and GitHub source code repositories used to generate the derivatives and tract labels in this benchmark. Each DOI references a specific, versioned instance of an App executed on the brainlife.io platform, ensuring transparency and full reproducibility of the preprocessing, tractography, and segmentation workflow.

|  |  |  |  |
| --- | --- | --- | --- |
| Application Name | Name | Brainlife DOI | Github Repository |
| FSL Anat (T1) | A273 | [10.25663/brainlife.app.273](https://doi.org/10.25663/brainlife.app.273) | https://github.com/brainlife/app-fsl-anat/tree/v1.1 |
| Align T1 to ACPC Plane (HCP-based) | A99 | [10.25663/bl.app.99](https://github.com/brainlife/app-hcp-acpc-alignme) | https://github.com/brainlife/app-hcp-acpc-alignme/tree/1.4 |
| Tissue-type segmentation | A239 | [10.25663/brainlife.app.239](https://github.com/brainlife/app-mrtrix3-5tt) | https://github.com/brainlife/app-mrtrix3-5tt/tree/binarize-v1.0 |
| Freesurfer | A0 | [10.25663/bl.app.0](https://github.com/brainlife/app-freesurfer) | https://github.com/brainlife/app-freesurfer/tree/1.12 |
| mrtrix3 preprocess | A68 | [10.25663/bl.app.68](https://github.com/brainlife/app-mrtrix3-preproc) | https://github.com/brainlife/app-mrtrix3-preproc/tree/1.8 |
| FSL Brain Extraction (BET) on DWI | A163 | [10.25663/brainlife.app.163](https://github.com/brainlife/app-FSLBET) | https://github.com/brainlife/app-FSLBET/tree/dwi |
| Fit Constrained Deconvolution Model for Tracking | A238 | [10.25663/brainlife.app.238](https://github.com/bacaron/app-mrtrix3-act) | https://github.com/bacaron/app-mrtrix3-act/tree/csd_generation-v1.0 |
| mrtrix3 - WMC Anatomically Constrained Tractography (ACT) | A319 | [10.25663/brainlife.app.319](https://github.com/brainlife/app-mrtrix3-act) | https://github.com/brainlife/app-mrtrix3-act/tree/1.4 |
| Anatomically Constrained Tractography using precomputed 5tt & CSD | A297 | [10.25663/brainlife.app.297](https://github.com/bacaron/app-mrtrix3-act) | https://github.com/bacaron/app-mrtrix3-act/tree/1.3 |
| White Matter Anatomy Segmentation | A188 | [10.25663/brainlife.app.188](https://github.com/brainlife/app-wmaSeg) | https://github.com/brainlife/app-wmaSeg/tree/3.9 |
| Remove Tract Outliers | A195 | [10.25663/brainlife.app.195](https://github.com/brainlife/app-removeTractOutliers) | https://github.com/brainlife/app-removeTractOutliers/tree/1.4 |
| app-tcks2tractmasks | A852 | [10.25663/brainlife.app.852](https://github.com/Junebeomstics/app-tcks2tractmasks) | https://github.com/Junebeomstics/app-tcks2tractmasks/tree/v1.2 |
| FOD (CSD) Registration | A849 | [10.25663/brainlife.app.849](https://github.com/gamorosino/app-registration-FOD) | https://github.com/gamorosino/app-registration-FOD/tree/main |

These three datasets present distinct challenges for deep learning tract segmentation, arising from systematic differences in acquisition properties and population characteristics. The datasets differ in diffusion shell composition and voxel resolution (**Fig. 2a**): more shells enable denser angular sampling of the diffusion signal, and higher resolution allows fODF peaks to encode finer spatial detail, each contributing distinct aspects of the information available for model learning and the quality of predicted tract masks. The datasets also span markedly different age distributions, representing pediatric brain development, adult maturity, and aging-associated white matter changes (**Fig. 2b**). As shown in **Fig. 2c**, HCP_1057_ subjects yield more complete arcuate fasciculus tract masks and streamlines than subjects from Cam-CAN_288_ and PING_108_, and this difference is amplified for subjects with atypical data quality. Together, this benchmark allows us to assess how robustly models trained on one acquisition protocol and population generalize when applied to data with substantially different characteristic (see **Methods** for dataset details).

**Figure 2.**
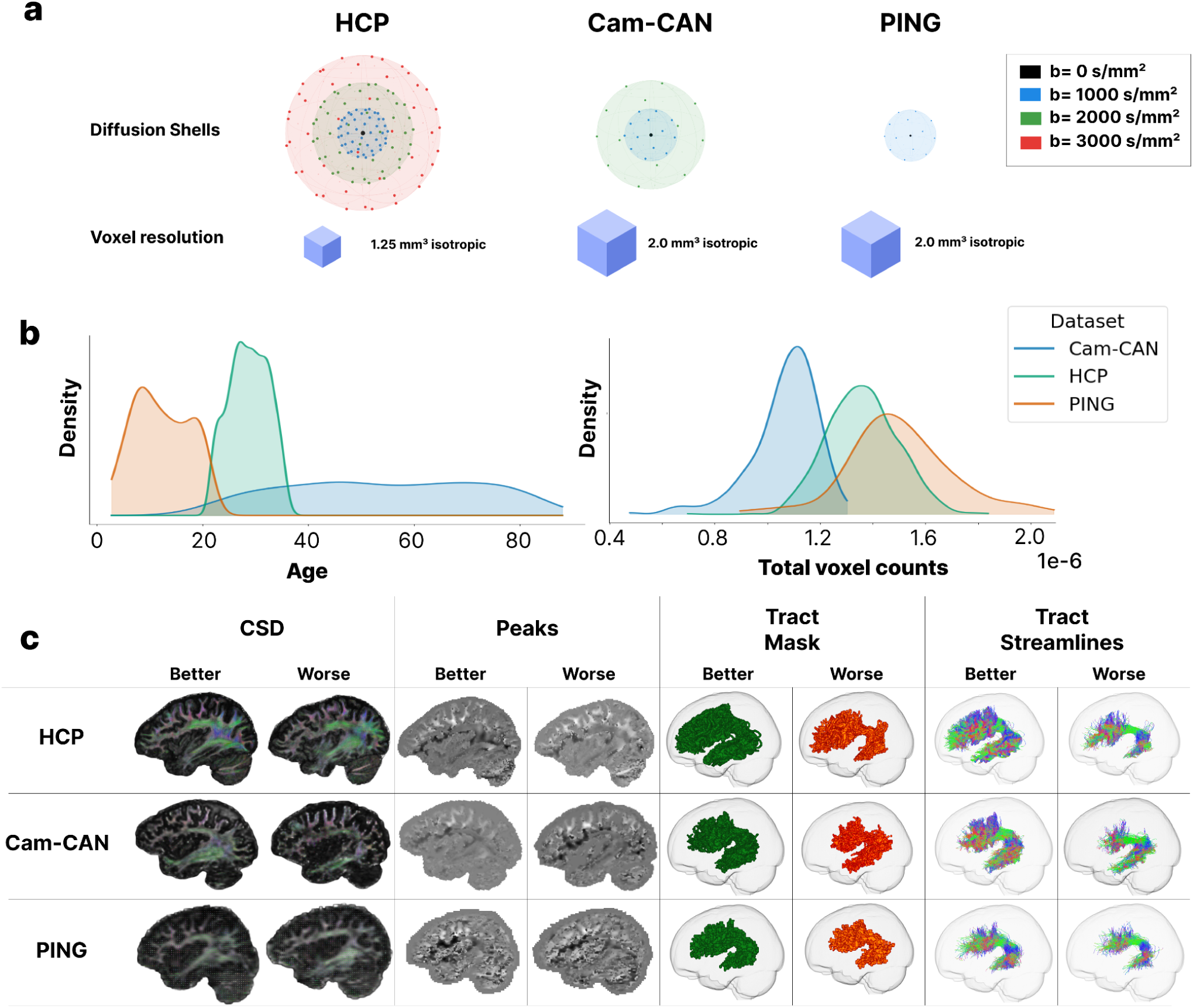
Acquisition parameters, data quality variability, and demographic composition of the three-dataset benchmark. **a.** Acquisition parameters across the three datasets (HCP, Cam-CAN, and PING). Diffusion shell compositions are visualized in q-space, where each b-value shell is represented as a sphere and individual b-vectors (gradient directions) as points distributed on its surface (b-values color-coded: b = 0 s/mm² black, b = 1000 s/mm² blue, b = 2000 s/mm² green, b = 3000 s/mm² red). Voxel resolution indicates the spatial detail of the diffusion MRI data; smaller voxels capture finer anatomical structures. **b,** Dataset distribution characteristics. Left: Age distributions across HCP_1057_ (green, 22–35 years), Cam-CAN_288_ (blue, 18–87 years), and PING_108_ (orange, 3–21 years). Right: Total brain voxel counts (×10⁶) within 61 tract masks. **c.** Data derivatives in the benchmark datasets. Each row shows two subjects from HCP_1057_, Cam-CAN_288_, and PING_108_, selected to represent typical (Good) or atypical (Bad) quality based on the voxel counts in the Arcuate Fasciculus tract mask. Columns show constrained spherical deconvolution (CSD) fiber orientation distributions (Lmax = 8, 6, 6), fiber orientation peaks (principal direction x component), binary tract masks for left arcuate fasciculus (green: typical quality; red: atypical quality), and tractography streamlines for left arcuate fasciculus (colored by local orientation).

### Overview of Models and Evaluation Framework

Below we perform a series of experiments to demonstrate the challenges that the individual datasets provide for different deep learning models. More specifically, we put emphasis on cross-dataset transfer and generalization, which was not possible with previous benchmark datasets, evaluating each model across 15 training–test combinations spanning HCP_1057_, Cam-CAN_288_, and PING_108_ (**Table 4**). In all experiments, models processed fODF peak directions estimated via Constrained Spherical Deconvolution ^64^ and predicted 61 three-dimensional binary masks ^52^ representing white matter tract volumes. Whereas ref ^28^ trained a U-Net on individual 2D slices of fODF peaks, all architectures in our benchmark were implemented as fully volumetric models to capture 3D spatial context (**Supplementary Fig. 1**).

**Table 4.** Experimental settings to evaluate the model performances in within-domain and out-of-domain generalization.

|  |  |
| --- | --- |
| **Train** | **Test** |
| **Within-Domain** |  |
| HCP<sub>1057</sub> | HCP<sub>1057</sub> |
| Cam-CAN<sub>288</sub> | Cam-CAN<sub>288</sub> |
| PING<sub>108</sub> | PING<sub>108</sub> |
| HCP<sub>1057</sub>&Cam-CAN<sub>288</sub>&PING<sub>108</sub> | HCP<sub>1057</sub> |
| HCP<sub>1057</sub>&Cam-CAN<sub>288</sub>&PING<sub>108</sub> | Cam-CAN |
| HCP<sub>1057</sub>&Cam-CAN<sub>288</sub>&PING<sub>108</sub> | PING |
| **Out-of-Domain** |  |
| HCP<sub>1057</sub> | Cam-CAN<sub>288</sub> |
| HCP<sub>1057</sub> | PING<sub>108</sub> |
| Cam-CAN<sub>288</sub> | PING<sub>108</sub> |
| Cam-CAN<sub>288</sub> | HCP<sub>1057</sub> |
| PING<sub>108</sub> | HCP<sub>1057</sub> |
| PING<sub>108</sub> | Cam-CAN<sub>288</sub> |
| HCP<sub>1057</sub>&PING<sub>108</sub> | Cam-CAN<sub>288</sub> |
| HCP<sub>1057</sub>&Cam-CAN<sub>288</sub> | PING<sub>108</sub> |
| Cam-CAN<sub>288</sub>&PING<sub>108</sub> | HCP<sub>1057</sub> |

We evaluated four deep learning architectures spanning from purely convolutional to transformer-based approaches (**Supplementary Fig. 2**). More specifically, we evaluated TractSeg ^28,29^, a U-Net-based architecture that established the current standard for voxel-based WMTS. We then tested MedNeXt ^53^, which consists of ConvNeXt-based blocks with depthwise convolutions. To assess an architecture capable of capturing more global relationships across fODF peaks, we included SwinUNETR ^54^, which combines local convolutional layers with hierarchical self-attention mechanisms for modeling long-range dependencies. Finally, we tested whether pretrained foundation models for general image segmentation tasks can be applied to WMTS. Specifically, we finetuned the pretrained Segment Anything Model^65^ to optimize for WMTS, following the strategies proposed for volumetric medical segmentation tasks, called Modality-Agnostic-SAM ^55^. We quantified segmentation accuracy using the Dice coefficient, the standard metric for WMTS evaluation ^44,46^. The Dice coefficient measures spatial overlap between predicted and ground-truth tract masks as follows:

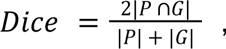

where P and G represent the predicted and ground-truth segmentation.

### Performance benchmarking under within- and out-of-domain settings

We evaluated model performance under two experimental regimes: *within-domain* and *out-of-domain*. In within-domain experiments, models were trained and tested on the same dataset, so the held-out test set came from the same distribution the model had already seen during training. These analyses estimate the best performance a model can reach on a given dataset and have been reported previously ^28,29,32^, although not at the scale or heterogeneity considered here. In contrast, out-of-domain experiments tested generalization by evaluating models on datasets that were absent from training. This setting introduces a *domain shift*, a change in the underlying data distribution between training and testing that arises from differences in acquisition protocols and study populations. Such analyses remain rare, largely because inconsistent tract labeling across datasets has historically restricted evaluation to qualitative comparisons.

### Within-domain experiments

We evaluated model performance under within-domain settings, where training and testing data were sampled from the same source dataset (i.e., both drawn from HCP_1057_, Cam-CAN_288_, or PING_108_). Within this framework, we distinguished between two training strategies to assess the impact of dataset diversity on model generalization; *homogeneous training and heterogeneous training*. Under *homogeneous training*, models were trained exclusively on data from a single dataset and then evaluated on held-out test data from that same dataset (e.g., trained on HCP_1057_, tested on HCP_1057_). Under *heterogeneous training*, models were trained on a combined training dataset from all three sources (HCP_1057_, Cam-CAN_288_, and PING108), then evaluated on held-out test sets from each individual dataset (e.g., trained on HCP_1057_&Cam-CAN_288_&PING108, and tested on HCP_1057_). This heterogeneous approach tests whether exposure to greater population diversity during training enhances model performance on each target dataset compared to training on that dataset alone. All experiments used a 70-15-15 split for training, validation, and testing, respectively.

#### Homogeneous training results

When testing using homogeneous training conditions, models achieved the highest accuracy on the HCP1057 dataset (Dice coefficient across models: 0.7919 ± 0.0193), followed by the PING108 dataset (Dice coefficient across models: 0.7356 ± 0.0250), and the Cam-CAN_288_ dataset (Dice coefficient across models: 0.6742 ± 0.0401; see **Fig. 3a**). Statistical significance across experiments was established using a two-way ANOVA with model (TractSeg, MedNeXt, SwinUNETR, MA-SAM) and experimental condition (homogeneous HCP1057, homogeneous Cam-CAN_288_, homogeneous PING108, heterogeneous to HCP1057, heterogeneous to Cam-CAN_288_ heterogeneous to PING108) as factors. The analysis revealed significant main effects of both model (F(3,1736) = 363.21, *p < 0*.001, η² = 0.124) and experimental condition (F(5,1736) = 1172.68, *p < 0*.001, η² = 0.67), as well as a significant model × condition interaction (F(15,1736) = 6.49, *p < 0*.001, η² = 0.01). These results indicate that model performance varied significantly across architectures and experimental conditions, with the interaction suggesting that relative model rankings depended on the specific training-testing configuration.

**Figure 3.**
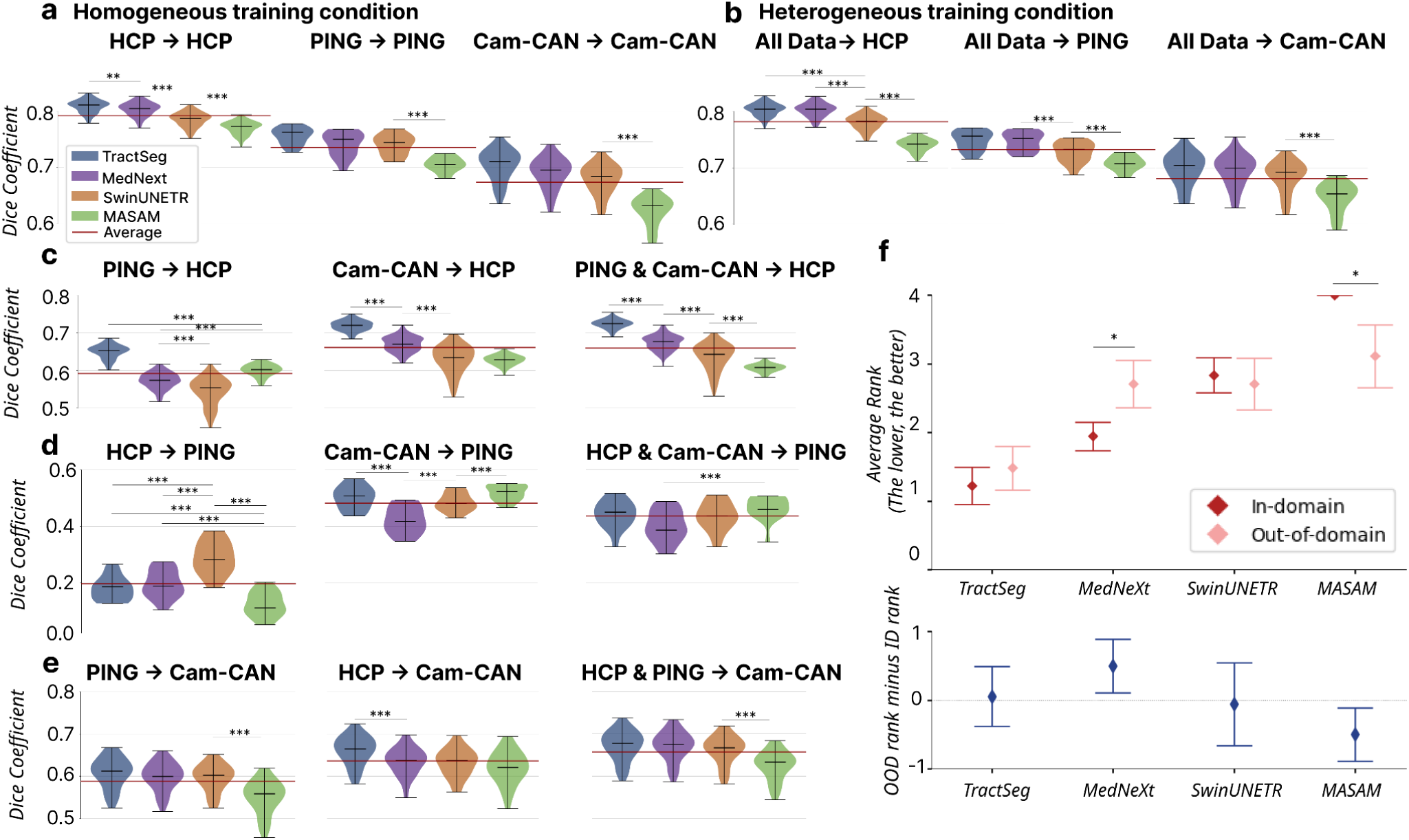
Within-domain and out-of-domain performance across models and datasets. Violin plots show Dice coefficient distributions across 61 white matter tracts (WMTs) with 95% confidence intervals. Red horizontal lines indicate the mean Dice coefficient across models. Dataset subscripts denoting sample size (HCP_1057_, Cam-CAN_288_, PING_108_) are omitted from panel labels for brevity. **a,** Within-domain performance under homogeneous training. Models were trained and tested on the same dataset. **b,** Within-domain performance under heterogeneous training: models trained on all three datasets, tested on one. **c-e,** Out-of-domain generalization. Models were trained on one or two source datasets and tested on the remaining held-out dataset: HCP_1057_ (c), PING_108_ (d), or Cam-CAN_288_ (e). **f,** Model ranking analysis. Top: average ranks (1=best, 4=worst) for within-domain (dark red) and out-of-domain (light pink) conditions. Bottom: rank differences (out-of-domain minus within-domain), where negative values indicate better out-of-domain performance.

Given the significant interaction effect, we performed post-hoc pairwise t-tests with Bonferroni correction to examine specific model differences within each condition. Under homogeneous training conditions, models consistently ranked in the following performance order: TractSeg, MedNeXt, SwinUNETR, and MA-SAM. On HCP_1057_, all pairwise model comparisons showed significant differences in performance (*p < 0*.001). In contrast, on PING_108_ and Cam-CAN_288_, only MA-SAM differed significantly from the other three models (*p* < 0.001), while TractSeg, MedNeXt, and SwinUNETR performed comparably. Overall, performance was highest on HCP_1057_ and lowest on Cam-CAN_288_, a pattern that likely reflects systematic differences in diffusion shell composition, voxel resolution, and population age across datasets.

#### Heterogeneous training results

Training with more heterogeneous datasets resulted in lower overall model performance (**Fig. 3b**; compared to homogeneous training, see **Fig. 3a**). Specifically, when evaluated on the HCP1057 dataset, TractSeg (t = 4.66, *p < 0*.001), SwinUNETR (t = 3.15, *p < 0*.001), and MA-SAM (t = 23.19, Bonferroni-corrected p < 0.001) exhibited significant performance decreases under heterogeneous training (MedNeXt did not show a significant difference). When evaluated on the PING108 dataset, no significant differences were observed across any models between homogeneous and heterogeneous training. Finally, when evaluated on the Cam-CAN_288_ dataset, only MA-SAM showed a significant difference (t = -4.2, *p < 0*.001), with heterogeneous training slightly improving its performance.

### Out-of-domain experiments

We designed out-of-domain experiments to assess how well models generalize across datasets with distinct acquisition protocols and population characteristics. We also tested whether combining source datasets provides complementary performance gains. Together, these comparisons reveal the distributional similarity among the three cohorts. We trained each model on one dataset and tested it on the remaining two, yielding six single-dataset training scenarios (**Fig. 3c-e**). Additionally, we trained models on two datasets combined, and tested them on the third, creating three two-dataset training scenarios (**Fig. 3c-e**).

#### Generalization to densely sampled adult data (HCP_1057_)

First, we trained models jointly or separately on Cam-CAN_288_ and PING_108_, then tested them on the 159 HCP_1057_ test subjects (**Fig. 3c**). This comparison tested whether models trained on sparser acquisitions with broader age ranges can generalize to the most densely sampled data in the benchmark, and whether combining two source datasets improves transfer over using either alone. A two-way ANOVA revealed significant main effects for both model architecture (F(3, 1896) = 1010.59, p < 0.001, η² = 0.399) and training setting (F(2, 1896) = 1093.68, p < 0.001, η² = 0.288), with a significant interaction (F(6, 1896) = 79.03, *p* < 0.001, η² = 0.062), indicating that model rankings varied depending on the source training data. In PING_108_→HCP_1057_, models ranked in the order TractSeg, MA-SAM, MedNeXt, and SwinUNETR, with all pairwise comparisons reaching significance (*p* < 0.001). In Cam-CAN_288_→HCP_1057_, models followed the same ranking pattern as the homogeneous HCP_1057_ condition (TractSeg, MedNeXt, SwinUNETR, MA-SAM), with all pairwise differences being significant (p < 0.001). Models trained on both datasets combined (PING_108_ & Cam-CAN_288_→HCP_1057_) followed this same ranking, with all pairwise differences being significant (p < 0.001). Across training settings, Cam-CAN_288_-trained models (Dice: 0.661 ± 0.044) consistently outperformed PING_108_-trained models (Dice: 0.592 ± 0.046). Additionally, adding PING_108_ to Cam-CAN_288_ training provided no additional benefit (Dice: 0.660 ± 0.048), suggesting that Cam-CAN_288_’s multi-shell protocol and broader age coverage already capture many of the features relevant for predicting HCP_1057_ anatomy. Nevertheless, even the best-performing transfer condition (Cam-CAN_288_→HCP_1057_, Dice: 0.661± 0.044) fell substantially below the homogeneous HCP_1057_ baseline (Dice: 0.7919 ± 0.0193), confirming that generalization from sparser to denser acquisitions remains difficult for all architectures.

#### Generalization to the developing brain (PING_108_)

Next, we trained models jointly or separately on HCP1057 and Cam-CAN_288_, then tested them on the 17 PING108 test subjects (**Fig. 3d**). This comparison tested whether diffusion shell similarity and population age overlap predict transfer success to developing brains. Cam-CAN_288_ and PING_108_ share comparably sparse shell coverage, whereas HCP_1057_’s dense three-shell protocol diverges from PING_108_’s single-shell acquisition. In terms of age, both Cam-CAN288’s lifespan range and HCP1057’s young-adult range overlap only marginally with PING_108_’s pediatric population. A two-way ANOVA revealed significant main effects for both model architecture (F(3, 192) = 10.49, *p* < 0.001, η² = 0.028) and training setting (F(2, 192) = 415.53, *p* < 0.001, η² = 0.730), with a significant interaction (F(6, 192) = 13.95, *p* < 0.001, η² = 0.074), indicating that model rankings varied depending on the source training data. In HCP_1057_→PING_108_, SwinUNETR significantly outperformed other models (p < 0.001). In Cam-CAN_288_→PING_108_, models ranked in the order MA-SAM, TractSeg, SwinUNETR, and MedNeXt, with MA-SAM significantly outperforming SwinUNETR (p < 0.001) and MedNeXt significantly underperformed compared to other models (p < 0.001). Models trained on both datasets combined (HCP_1057_ & Cam-CAN_288_→PING_108_) showed MA-SAM significantly outperforming MedNeXt (*p* < 0.001). Across training settings, Cam-CAN_288_-trained models (Dice: 0.4813 ± 0.0519) substantially outperformed HCP_1057_-trained models (Dice: 0.1964 ± 0.0759; p < 0.001 for all models), and combining the two datasets yielded only intermediate performance (Dice: 0.4349 ± 0.0528; Fig. 3d, middle row), suggesting that adding HCP_1057_ to Cam-CAN_288_ training degraded rather than improved transfer to pediatric data. Even the best transfer condition (Cam-CAN_288_→PING_108_, Dice: 0.4813 ± 0.0519) fell far below the homogeneous PING_108_ baseline (Dice: 0.7356 ± 0.0250). The steepest drop occurred in HCP_1057_→PING_108_ suggests that mismatched shell coverage and limited age overlap jointly imposed a severe constraint on generalization to developing brains.

#### Generalization to the aging lifespan cohort (Cam-CAN_288_)

Last, We trained models on HCP1057 and PING108, then tested them on 44 Cam-CAN_288_ test subjects (**Fig. 3e**). A two-way ANOVA revealed significant main effects for both model architecture (F(3, 516) = 19.82, *p < 0*.001, η² = 0.081) and training setting (F(2, 516) = 75.71, *p < 0*.001, η² = 0.206). In the PING→Cam-CAN_288_ setting, only the differences between MA-SAM and the other three models reached statistical significance (Bonferroni-corrected *p < 0*.001). On the other hand, when trained exclusively on HCP1057 data (HCP1057→Cam-CAN_288_), TractSeg substantially outperformed all other architectures (p < 0.001 for all pairwise comparisons). HCP1057-trained models (Dice: 0.6365 ± 0.0374) consistently outperformed PING108-trained models (dice: 0.5884 ± 0.0413) across all architectures. Training jointly on both datasets improved Cam-CAN_288_ prediction performance compared to training on them separately (Dice: 0.6573 ± 0.0387). This result suggests that HCP1057 and PING108 provide complementary information for predicting Cam-CAN_288_ dataset.

#### Architecture rankings shift under domain shift

Next, we sought to identify which architectures perform best across varying dataset conditions (**Fig. 3f**). To do so, we ranked each model’s performance within each experimental setting, then averaged these ranks separately for within-domain and out-of-domain scenarios. Across all within-domain conditions, TractSeg ranked highest on average (mean rank: 1.22), followed by MedNeXt (1.94), SwinUNETR (2.83), and MA-SAM (4.0). In out-of-domain settings, TractSeg maintained the top position (mean rank: 1.48), but the relative performance of other models shifted substantially: MedNeXt dropped in relative ranking (1.94→2.70; rank difference: +0.76), while transformer-based models improved, with SwinUNETR (2.83→2.70; −0.13) and MA-SAM (4.0→3.11; −0.89) both performing better out-of-domain than within-domain (**Fig. 3f**, lower panel). These results indicate that while CNN-based models achieve superior within-domain performance, transformer-based architectures show stronger generalization under domain shift, narrowing or reversing the performance gap observed in within-domain evaluation.

#### The benefit of volumetric processing is architecture-dependent

We next examined how volumetric processing, tract morphology, and tract volume each affect segmentation performance across architectures and datasets. The experiments are reported as **Supplementary Results 3-6**. We examined whether volumetric processing contributed to these performance differences (**Supplementary Result 3**). Volume-based TractSeg significantly outperformed its slice-based counterpart in 6 of 15 settings and underperformed in only 1, whereas volume-based and slice-based SwinUNETR showed no significant difference in 10 of 15 settings, with slice-based SwinUNETR outperforming in three out-of-domain transfers. The performance benefit of volumetric processing is therefore architecture-dependent: TractSeg exploits 3D context more consistently across settings than SwinUNETR.

#### Tract volume predicts segmentation accuracy

We also evaluated how segmentation accuracy varied across individual tracts (**Supplementary Result 4**). To determine whether tract volume underlies this variability, we correlated voxel count with Dice coefficient for 3D TractSeg across three within-domain settings (**Supplementary Result 5**). Larger tracts consistently achieved higher Dice scores across all three datasets, with moderate positive correlations: r = 0.50 (p = 3.96 × 10⁻⁵) for HCP_1057_→HCP_1057_, r = 0.52 (p = 1.54 × 10⁻⁵) for Cam-CAN_288_→Cam-CAN_288_, and r = 0.56 (p = 3.07 × 10⁻⁶) for PING_108_→PING_108_.

#### Ensemble integration partially offsets degradation under severe domain shift

Because no single architecture generalized reliably across all transfer scenarios, we asked whether integrating multiple models could compensate for individual failures under domain shift. We therefore introduce Ensemble White Matter Tract Segmentation (EWMTS), which combines predictions from five models using two consensus strategies, majority voting and STAPLE, across all 15 experimental settings (see **Supplementary Result 7** and **Methods**). In within-domain experiments, majority voting performed between the median and best individual model, whereas STAPLE consistently underperformed compared to worst models. In out-of-domain transfers, the ranking reversed: STAPLE outperformed majority voting by 67% in the most challenging scenario (HCP_1057_→PING_108_; Dice: 0.30 vs. 0.18, *p* < 0.001) and showed a similar advantage in PING_108_→HCP_1057_ (STAPLE: 0.61 vs. majority voting: 0.58, *p* < 0.001). Threshold sensitivity analysis further showed that STAPLE maintained stable performance across binarization thresholds, whereas majority voting degraded steeply above the optimal threshold in out-of-domain settings (**Supplementary Result 8**). Qualitative comparisons of predicted tract masks showed that STAPLE better preserved tract coverage under severe domain shift, particularly for smaller association tracts (**Supplementary Result 9**).

## DISCUSSION

This study introduces the first benchmark dataset designed to evaluate cross-domain generalization in white matter tract segmentation, spanning pediatric, adult, and aging cohorts acquired on three different scanners. Applying four state-of-the-art architectures to this benchmark, we found that all models suffer substantial performance degradation under domain shift, most severely when adult-trained models are applied to pediatric data (Dice: 0.196). Such degradation is invisible under conventional within-domain evaluation, where models appear to perform well despite failing to generalize. By openly releasing this dataset and evaluation framework, we provide a foundation for the community to develop, benchmark, and refine segmentation models under the heterogeneity of real-world neuroimaging data.

Our benchmark addresses a critical limitation of existing evaluation practice. Most prior studies trained models on a single manually curated dataset of 105 HCP subjects (HCP_105_) and assessed generalization either qualitatively or on a small subset of manually labeled tracts, making it difficult to test whether models generalize reliably to new datasets. This study leverages brainlife.io to implement a fully traceable preprocessing pipeline alongside a benchmark of 1,455 subjects spanning three public datasets, capturing realistic variability in scanner hardware, acquisition protocols, and population demographics that the HCP_105_ data cannot represent. Researchers can either adopt this benchmark directly or apply the identical pipeline to their own data, enabling consistent cross-study comparisons and incremental expansion as new cohorts become available.

Within-domain performance in homogeneous conditions reflects the joint influence of acquisition quality and population heterogeneity on segmentation accuracy (**Fig 3a**). HCP_1057_ yields the highest accuracy across all architectures, consistent with its narrow young-adult age range and three-shell, high-resolution acquisition, which together provide anatomically homogeneous data with information-rich fODF peaks. In contrast, PING_108_ and Cam-CAN_288_ yield lower accuracy. Both datasets combine sparser angular sampling with greater inter-subject anatomical variability, arising from active neurodevelopment in PING_108_ and aging-related white matter changes in Cam-CAN_288_. Within these more challenging datasets, the performance gap is further amplified for smaller tracts: large, well-defined tracts such as the SLF I/II and corpus callosum segments maintain Dice coefficients above 0.80 across all three datasets, whereas smaller tracts such as the uncinate fasciculus and posterior arcuate achieve comparable accuracy only in HCP_1057_, with markedly lower and more variable performance in Cam-CAN_288_ and PING_108_ (**Supplementary Fig. 4–5**). This dissociation confirms that intrinsic data properties set a performance ceiling for fine anatomical structures that architectural advances alone cannot overcome.

Heterogeneous training, in which models were trained on all three datasets combined, tested whether greater data diversity would improve generalization by covering a broader range of anatomy and acquisition protocols. However, the results indicate that dataset compatibility, not sample size alone, determines transfer performance. Within a single dataset, expanding HCP training data from 105 to 1,057 subjects improved accuracy across all architectures (**Supplementary Fig. 6**), confirming that sample size matters when data are drawn from a consistent distribution. However, pooling all three datasets during training provided no benefit over training on a single matched dataset, and in some cases degraded performance (**Fig. 3a vs. 3b**). This pattern indicates that combining distributionally dissimilar datasets does not provide additive training benefit; rather, the model must simultaneously learn distinct feature patterns from each dataset, which reduces its specialization for any single target domain. Without explicit domain alignment, training on distribution-matched data therefore outperforms naive data aggregation.

Out-of-domain transfer performance varies systematically with the distributional distance between source and target datasets (**Fig. 3c**), a distance jointly shaped by diffusion shell composition and population characteristics. HCP_1057_ and Cam-CAN_288_ are the most similar pair in the benchmark: both offer multi-shell protocols and adult brain data, and bidirectional transfer between them yields moderate accuracy (Cam-CAN_288_→HCP_1057_, Dice: 0.661; HCP_1057_→Cam-CAN_288_, Dice: 0.637). PING_108_ diverges from both datasets, but more from HCP_1057_, whose dense three-shell protocol and narrow young-adult age range differ from PING_108_’s single-shell, pediatric data.

This simultaneous mismatch in both shell coverage and population age produces the most severe transfer failure in the benchmark (HCP_1057_→PING_108_, Dice: 0.196). In contrast, Cam-CAN_288_→PING_108_, yields substantially higher performance (Dice: 0.481), consistent with Cam-CAN_288_’s sparser shell coverage and partially overlapping age range with PING_108_. Jointly training on HCP_1057_ and PING_108_ partially improved Cam-CAN_288_ prediction, suggesting that HCP_1057_’s multi-shell protocol and PING_108’_’s broader age coverage each contribute features relevant to Cam-CAN_288_’s intermediate acquisition and demographic characteristics. These results carry a practical implication for model deployment: when the target cohort differs from available training data, selecting source datasets that match on both shell composition and population age will yield better transfer than defaulting to the largest or highest-resolution dataset available.

Across all within-domain conditions, CNN-based models consistently outperformed transformer-based architectures (TractSeg mean rank: 1.2, MedNeXt: 1.9, SwinUNETR: 2.8, MA-SAM: 4.0), a ranking that reflects the local spatial structure of fODF peak data (**Fig. 3e**). fODF peaks encode dominant fiber orientations within each voxel, a representation in which spatially local relationships carry the most discriminative information for tract boundary delineation. CNN-based models exploit this structure effectively through hierarchical local receptive fields, whereas transformer-based architectures compute global dependencies from early layers, a capacity that offers limited additional benefit when the informative signal is inherently local. Transformers also lack the translation equivariance built into CNNs, requiring substantially more training data to learn comparable spatial representation ^66,67^. To mitigate this limitation, we evaluated SwinUNETR, a hybrid architecture that combines local convolutions with global self-attention, and MA-SAM, which leverages large-scale pretraining on natural images. Nonetheless, neither strategy overcame the within-domain advantage of purely convolutional models, suggesting that the sample sizes currently available for tract segmentation are insufficient to realize the representational benefits transformers offer at scale.

Under domain shift, however, this ranking partially reversed: SwinUNETR and MA-SAM showed relative improvement in out-of-domain settings (**Fig. 3e**), suggesting that global attention, by not committing to local signal structure, tolerates changes in feature distributions more readily across acquisition protocols. MA-SAM nonetheless remained the weakest individual model across all conditions, indicating that large-scale pretraining on natural RGB images does not compensate for the domain gap to nine-channel fODF peak data; accordingly, it was excluded from ensemble experiments. Combining the remaining models through STAPLE ensemble integration partially mitigated the severe performance degradation observed in the most challenging transfers (e.g., HCP_1057_→PING_108_: STAPLE Dice 0.30 vs. best individual model Dice 0.18). These results suggest that when no single architecture generalizes reliably across all deployment scenarios, ensemble integration offers a practical strategy for maintaining segmentation accuracy under domain shift.

This study has three limitations. First, the three datasets differ simultaneously across multiple dimensions, including diffusion shell composition, voxel resolution, scanner hardware, and population age range, which prevents clean attribution of generalization failures to any single factor. Datasets that vary systematically along one dimension, such as multisite studies with harmonized protocols across age groups, would help disentangle these contributions. Second, all models were trained on fODF peak directions. Whether alternative input representations, such as full fiber orientation distributions, diffusion tensor scalars, or T1-weighted structural images, would alter within- and out-of-domain patterns remains untested. Third, our benchmark addresses only voxel-based segmentation; extending this evaluation framework to streamline-based methods would provide a more complete picture of where generalization succeeds and fails across the broader landscape of tract segmentation. Despite these limitations, the reproducible benchmark, cross-dataset evaluation framework, and ensemble strategy established here provide a foundation for developing segmentation tools that generalize reliably across diverse populations and acquisition protocols.

## ONLINE METHODS

### Data Sources

We used diffusion-weighted MRI (DWI) and anatomical MRI (aMRI) scans from participants in the Human Connectome Project Young Adults (HCP-YA) ^9^, Cambridge Centre for Ageing and Neuroscience (Cam-CAN) ^59^, and Pediatric Imaging, Neurocognition, and Genetics (PING)^60^ data to examine the segmentation performances of each model. The characteristics and sample sizes used for each dataset are as **Table 2**.

### Data Preprocessing Pipeline

#### brainlife.io

We conducted all preprocessing using the brainlife.io ^52^, a cloud-based neuroimaging data processing platform that enables reproducible and standardized analysis workflows. *brainlife.io* provides a comprehensive ecosystem that seamlessly integrates established neuroimaging toolkits, including MRtrix3, FSL, FreeSurfer, and other widely used software packages through containerized processing pipelines. A key advantage of the brainlife.io platform is its built-in provenance tracking system, which automatically records detailed metadata about each processing step, including the specific Apps used, parameter settings, and data transformations applied. This comprehensive provenance documentation ensures the full reproducibility of our preprocessing pipeline and enables transparent verification of all processing steps, making our tract segmentation model training reproducible. The Apps used are listed in **Table 3**.

#### Anatomical MRI (aMRI) Processing

All three datasets underwent preprocessing based on the HCP1057 Preprocessing Pipeline^1^. For HCP1057 data, we used preprocessed T1-weighted images (T1w_acpc_dc_restore_1.25.nii.gz), where “acpc_dc_restore” indicates that the images had already undergone ACPC alignment, distortion correction, and bias field correction with 1.25mm isotropic resolution. For Cam-CAN_288_ datasets, we performed bias correction and alignment to the anterior commissure-posterior commissure (ACPC) plane using the App [A273](https://doi.org/10.25663/brainlife.app.273) on *brainlife.io*. For PING108 data, we performed ACPC alignment without bias correction using App [A99](https://doi.org/10.25663/bl.app.99) on *brainlife.io*.

We generated gray-white matter interface masks for white matter tractography by segmenting T1-weighted images into tissue types using MRTrix3 ^68^ (App [A239](https://doi.org/10.25663/brainlife.app.239)). These masks served as seed regions for tractography analysis. Additionally, we processed T1-weighted images for cortical surface reconstruction using FreeSurfer’s recon-all pipeline ^69^ (App [A0](https://brainlife.io/app/58c56d92e13a50849b258801)).

#### Diffusion-weighted MRI (DWI) Processing & Fiber Modeling

For the HCP1057 data, we used minimally-preprocessed DWI images ^70^. For Cam-CAN_288_ and PING108 datasets, we preprocessed DWI images following the protocol in QSIPrep using App [A68](https://brainlife.io/app/5a813e52dc4031003b8b36f9) ^71^. Specifically, we denoised DWI images, removed Gibbs ringing using MRTrix3 ^68^ and corrected for susceptibility, motion, and eddy distortions using FSL’s *topup* and *eddy* functions ^72,73^. We applied eddy-current and motion correction via *eddy_cuda8.0* with outlier slice replacement (*repol*) from FSL ^74^. We used MRTrix3’s *dwigradcheck* functionality to check and correct misaligned gradient vectors following *top-up* and *eddy* ^68^. We debiased DWI images using ANT’s *n4* functionality ^75^ and cleaned background noise using MrTrix3.0’s *dwidenoise* ^76^. We registered preprocessed DWI images to structural (T1w) images using FSL’s *epi_reg* ^77^. We generated brain masks for DWI data using FSL’s *bet* ^73^ implemented as App [A163](https://brainlife.io/app/5c63a9c3f2362b00318046c2).

#### Constrained spherical deconvolution (CSD) model fitting

We fit the constrained spherical deconvolution (CSD) model ^78^ to preprocessed DWI data across different spherical harmonic orders (*L_max_*= 2,4,6,8 for HCP1057 and PING108 data, *Lmax* = 2,4,6 for Cam-CAN_288_ data) using *MRTrix* ^68^ implemented as App [A238](https://brainlife.io/app/5db224e98aeeee0f87f25c1f). We conducted this process within the tractography stage for PING108 data, using App [A319](https://brainlife.io/app/5db224e98aeeee0f87f25c1f).

#### Peak extraction and preprocessing

We extracted Peaks from the fitted CSD model to represent the three primary diffusion directions within each voxel. The extracted Peaks consisted of peak directions (unit vectors) and corresponding amplitudes, providing a discrete representation of underlying fiber orientations. For each peak, we obtained three spatial coordinates (x, y, z), resulting in a total of nine volumes when considering up to three Peaks per voxel (three Peaks × three coordinates = nine volumes). We replaced the NaN values in the Peaks with zero values. The Peaks reconstruction used *Lmax* =8 for the HCP1057 and PING108 datasets, while *Lmax* =6 was applied for the Cam-CAN_288_ dataset. These peak data were used as input to the model.

#### Tractography

We used Peaks and gray-white matter interface masks to perform anatomically-constrained probabilistic tractography (ACT) ^79^ using *MRTrix3* ^68^ implemented as App [A297](https://brainlife.io/app/5e88b8bf952fef39d57abcf2) or [A319](https://brainlife.io/app/58c56d92e13a50849b258801). We used *Lmax* =8 for HCP_1057_ and PING_108_ datasets, and *Lmax* =6 for Cam-CAN_288_ dataset. We generated 3 million streamlines for all datasets using a step size of 0.2 mm. For the HCP1057 and Cam-CAN_288_ datasets, we set the minimum and maximum streamline lengths to 25 and 250 mm, respectively, with a maximum curvature angle of 35^◦^. For the PING108 dataset, we set the minimum and maximum streamline lengths to 25 and 220 mm, respectively, with a maximum curvature angle of 35^◦^. We also generated diffusion tensor imaging (DTI) scalar maps, including fractional anisotropy (FA) and mean diffusivity (MD), within the same application.

#### White matter segmentation and cleaning

After generating a whole-brain tractography file, we used a modified version of the White Matter Query Language ^61^ to segment 61 WMT ^52,80^ automatically. This segmentation utilizes a combination of regions of interest (ROIs) from the FreeSurfer’s Destrieux 2009 parcellation^62^, as well as tract midpoint estimation, as inclusion and exclusion criteria to define a tract. The segmentation is implemented as an App https://brainlife.io/app/5cc73ef44ed9df00317f6288. Following the segmentation, streamline outlier methods were used ^13^, implemented in [App A195](https://brainlife.io/app/5cc9eca04b5e4502275edba6).

#### Binary Tract Mask Generation

Binary tract masks were generated using a script provided by TractSeg authors^28^. We implemented the script as App [A852](https://brainlife.io/app/67dc4fa6da629ed3b1e083d4). In a nutshell, for each tract, the App identifies the voxels through which the streamlines of a tract traverse; these voxels are set to 1, while all other voxels (those without tract streamlines) are set to 0. This method generates 61 WMT masks. These 61 tract-mask volumes are then combined into a single 4D file.

#### Registration to Standard Space

We transformed all non-HCP datasets into HCP1057 data space. To do so, we registered the FA images estimated from the CAM-CAN_288_ and PING108 datasets to an HCP1057 FA image registered to the MNI template^2^ ^28^. The affine registration map obtained through this process was then applied to the CSD data of the CAM-CAN_288_ and PING108 datasets (App [A849](https://brainlife.io/app/67d0b9371b629b467fa47511)). Peak images were then obtained for the two datasets using the standard-aligned CSD images. Finally, we applied the same registration to the tract masks generated in each dataset’s native space, ensuring that all data were aligned in the HCP1057 space.

### DL Model Training Strategy and Experimental Design

#### Data Preprocessing and Augmentation

All DL experiments were implemented using Peaks data as input, following the TractSeg project ^28^. For each WMT, the tract mask was used to represent the volume of the brain tissue touched by the WMT. The task of a DL Model was to segment the tract mask for a specific dataset and subject.

All Peaks and tract masks were preprocessed by cropping non-brain regions to achieve uniform dimensions of 144 x 144 x 144 from the original 145 x 174 x 145 resolution. We used zero-mean unit-variance normalization per channel to standardize Peaks intensity across subjects and acquisition parameters. During training, we applied data augmentation techniques to all samples ^29^. Specifically, Elastic deformation, Zooming, Gaussian Noise, and

Blur were used. These strategies increased the variability in the data, providing the DL Models with the ability to train on the datasets effectively.

#### Train and Validation Datasets

Multiple datasets were used in different experiments as the training and validation datasets. The HCP1057, Cam-CAN_288_, and PING108 datasets were used with a 70%-15%-15% split for training, validation, and testing, respectively. The 15% test data remained as held-out evaluation sets, while training and validation sets were randomly sampled from the remaining 85% of the data. All experiments were repeated three times to ensure statistical reliability. To evaluate model scalability and cross-domain adaptation capabilities, the following training-testing combinations were systematically evaluated, as shown in **Table 4**.

### Model Architectures and Implementation

#### TractSeg (Baseline CNN)

We implemented TractSeg based on the original authors’ codebase ^28,29^. Both 2D slice-based and 3D volumetric variants were evaluated to assess the impact of integrating spatial context. For 2D processing, individual slices were treated as separate samples within a batch configuration, maintaining the batch size of 47 as used in the original paper.

#### MedNeXt (Advanced CNN)

MedNeXt builds upon the self-configuring nnU-Net framework, incorporating ConvNeXt-inspired architectural enhancements, such as depth-wise separable convolutions and inverted bottleneck structures ^53^. The Base variant of MedNeXt was employed with 3 x 3 x 3 kernel sizes for 3D convolutions. To ensure a fair comparison across models, deep supervision was implemented using the same strategy employed in TractSeg. Only the 3D volumetric version was evaluated, processing complete brain volumes with a batch size of 2.

#### SwinUNETR (CNN-Transformer hybrid model)

We used SwinUNETR v2, which combines original Swin Transformer encoders with convolutional layers and U-Net-like decoder structures ^54^. Due to architectural constraints requiring specific input dimensions, zero-padding was applied to transform the 144 x 144 x 144 input volumes to 160 x 160 x 160 spatial dimensions. Both 2D slice-based and 3D volumetric implementations were evaluated to assess the impact of dimensionality on Transformer performance. Deep supervision was implemented consistently across all variants to maintain experimental parity. The model processes data with a batch size of 2 for volumetric processing, while slice-based processing utilizes individual slices as batch elements.

#### MA-SAM (Foundation Model)

The Modality-Agnostic Segment Anything Model (MA-SAM) is a foundation model approach for three-dimensional volumetric data, such as medical images^55^. MA-SAM is initialized with Vision Transformer weights pre-trained on 2D natural images from the original Segment Anything Model ^65^. To adapt the model for 3D medical volumes, additional 3D CNN layers were integrated with the original Transformer blocks to capture three-dimensional spatial information. The model employs Parameter-Efficient Transfer Learning (PETR) methods, enabling selective fine-tuning of the Transformer encoder while maintaining the beneficial properties of pre-trained features. This approach balances efficiency and accuracy by avoiding both complete model freezing and full end-to-end training. The ViT-Large variant was selected based on available computational resources. **Table 5** compares the efficiency of models used in our study based on the number of parameters, GFLOPs, and Throughput.

**Table 5.** Model Comparison: Parameters, Computational Cost, and Throughput.

|  |  |  |  |  |
| --- | --- | --- | --- | --- |
| **Model** | **Total Parameters** | **Trainable Param** | **GFLOPs** | **Throughput (Vol/sec)** |
| TractSeg 2D | 37.03M | 37.03M | 1008.79 | 304.74 |
| TractSeg 3D | 17.15M | 17.15M | 2574.56 | 11.98 |
| SwinUNETR 2D | 28.69M | 28.69M | 393.2 | 194.81 |
| SwinUNETR 3D | 18.37M | 18.37M | 445.41 | 4.152 |
| MASAM 3D (ViT<sub>large</sub>) | 375.01M | 70.90M | 5744.28 | 1.22 |
| MedNeXt 3D | 10.59M | 10.59M | 250.08 | 7.86 |

### Training Configuration and Optimization

We trained our models using consistent optimization strategies to ensure fair comparison ^29^. All models were trained using binary cross-entropy loss with sigmoid activation functions in the final layer to produce voxel-wise probability values (ranging from 0 to 1). For a given target *yi*, an output of the network *ŷ*_*i*_, where *i* identifies the number of classes, *N*, the loss is calculated using a log-likelihood function:

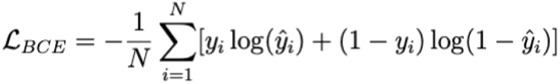

The AdamaX optimizer was employed for stable convergence across different architectures, coupled with Cosine Annealing Warm Restarts learning rate scheduling. We used binary cross-entropy loss to train the models. For batch configuration, 2D slice-based models processed individual slices as batch elements with single-channel input, while 3D volumetric models used a batch size of 2 due to memory constraints. We applied deep supervision techniques across TractSeg, MedNeXt, and SwinUNETR to improve gradient flow and improve convergence, with MA-SAM utilizing its inherent architectural design without requiring additional supervision.

### Ensemble approach

To evaluate ensemble-based tract segmentation, we aggregated binary predictions from five base models (TractSeg, TractSeg3D, SwinUNETR, SwinUNETR3D, and MedNeXt3D) across subjects using two fusion strategies: majority voting and STAPLE. For STAPLE fusion, we used the SimpleITK STAPLEImageFilter with a confidence weight of 1.0 and a maximum of 100 iterations. For majority voting, we averaged the predicted maps across models. Final ensemble masks were obtained by applying a binary threshold of 0.5 to the fused outputs.

## Acknowledgements

This research was supported by the following grants: Wellcome Trust (grant no. 226486/Z/22/Z, Principal Investigator F. Pestilli); NINDS UM1NS132207, BRAIN CONNECTS: Center for Mesoscale Connectomics (Principal Investigator K. Ugurbil); and NINDS U24NS140384, BRAIN CONNECTS: The Axonal Projectome EXchange (APEX) (Principal Investigator F. Pestilli). We thank Amazon Web Services Open Data Sponsorship Program for supporting data storage for brainlife.io

## Footnotes

1 https://github.com/Washington-University/HCPpipelines.git

2 https://github.com/MIC-DKFZ/TractSeg/blob/master/tractseg/resources/MNI_FA_template.nii.gz

