## Supplementary Material for "Benchmarking Generalizability in Deep Learning-Based White Matter Tract Segmentation"

#### Summary paragraph

White matter tracts (WMTs) are the brain's structural foundation for information transfer, underlying essential cognitive and behavioral functions. While diffusion MRI and tractography enable non-invasive mapping of these pathways, automated segmentation often lacks generalizability across diverse data sources. We conducted a systematic, cross-dataset evaluation of four state-of-the-art deep learning architectures, benchmarking their performance across independent datasets with varying acquisition protocols and populations. CNN-based models such as TractSeg achieved the highest within-domain accuracy, but performance dropped sharply under domain shift, most severely when we applied adult-trained models to pediatric data. To address this degradation, we introduce Ensemble White Matter Tract Segmentation (EWMTS), which combines complementary models to partially recover accuracy under domain shift, although performance still falls short of within-domain levels. By openly releasing this benchmark and a reproducible processing pipeline, we provide the neuroimaging community with a framework to develop and benchmark segmentation models across the heterogeneity of real-world neuroimaging data.

#### Author affiliations

[1] Department of Psychology, Center for Perceptual Systems, The University of Texas, Austin, TX 78712
[2] Department of Neuroscience, Center for Learning and Memory, The University of Texas, Austin, TX 78712

#### Competing interests.

The authors declare no competing financial interests.

#### Correspondence.

Franco Pestilli

#### Contribution.

J.B. and F.P. designed the experiments, with input from G.A. J.B. implemented the models, ran the benchmarking experiments, and performed the statistical analyses and visualization. G.A. contributed software tools for tract mask generation and registration. J.B. and F.P. wrote the first draft. J.B., G.A. and F.P. edited the manuscript.

#### Keywords

White Matter Tract Segmentation, Deep learning, Benchmark, Domain shift

#### Acknowledgements

This research was supported by the following grants: Wellcome Trust (grant no. 226486/Z/22/Z, Principal Investigator F. Pestilli); NINDS UM1NS132207, BRAIN CONNECTS: Center for Mesoscale Connectomics (Principal Investigator K. Ugurbil); and NINDS U24NS140384, BRAIN CONNECTS: The Axonal Projectome EXchange (APEX) (Principal Investigator F. Pestilli). We thank Amazon Web Services Open Data Sponsorship Program for supporting data storage for brainlife.io

### Supplementary Result 1: Comparison of Volume- and Slice-based approaches

White matter tract segmentation models differ fundamentally in how they integrate spatial information from diffusion MRI data. The original TractSeg pipeline adopted a slice-based approach [1]: Specifically, fODF peaks and tract masks at 144×144×144 resolution (cropped from 145×174×145 to remove non-brain background) were randomly sampled along the x-, y-, and z-axes, and a 2D U-Net was trained on these individual slices. During inference, predictions from all three orthogonal orientations were generated independently and then averaged to produce the final 3D segmentation mask. While this approach substantially reduces memory requirements, it discards the volumetric spatial context available across adjacent slices. In contrast, a volume-based approach processes the complete 3D fODF peak volume in a single forward pass through a 3D encoder-decoder, allowing the model to jointly optimize for spatial relationships across all three anatomical dimensions (**Supplementary Fig. 1**). All four architectures in the main benchmark were implemented as volume-based models. To isolate the contribution of volumetric context from architectural differences, we additionally evaluated 2D and 3D implementations of TractSeg and SwinUNETR across all 15 experimental conditions (**Supplementary Fig. 3**).

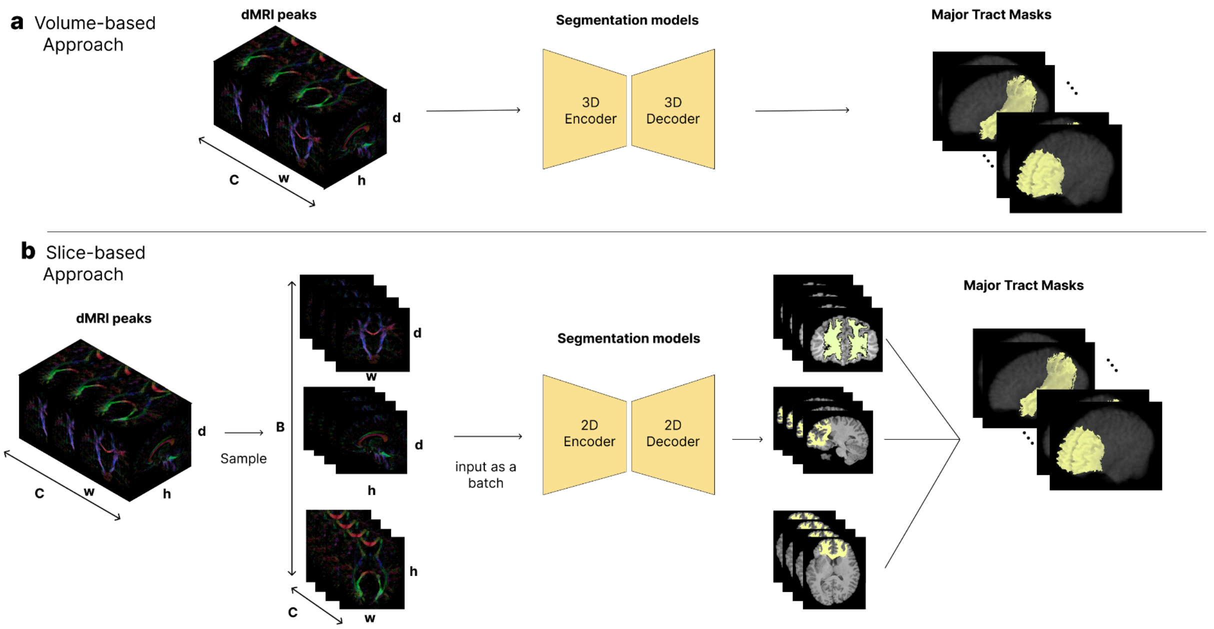


**Supplementary Figure 1.** Schematic comparison of volume-based and slice-based approaches for white matter tract segmentation. (a) Volume-based approach: the full 3D dMRI peak volume (dimensions C × W × H × D) is fed directly into a segmentation model with a 3D Encoder and 3D Decoder, which outputs 61 Major Tract Masks in a single forward pass. (b) Slice-based approach: During training stage, from the same 3D input volume, individual 2D slices are sampled along three anatomical orientations (axial, sagittal, and coronal) and processed as a batch through a 2D Encoder and 2D Decoder. During the inference stage, slices of peaks are used to predict Major Tract Masks and subsequently aggregated to reconstruct 3D tract masks.

**Supplementary Result 2: Detailed architectures of benchmark models**

The four architectures evaluated in this benchmark span a spectrum from established convolutional designs to transformer-based and foundation model approaches (**Supplementary Fig. 2**). TractSeg employs a standard 3D U-Net with 3×3×3 convolutions, batch normalization, and skip connections that preserve spatial detail across encoder and decoder pathways. Deep supervision, originally introduced in TractSeg [2], applies auxiliary losses at intermediate decoder scales to strengthen gradient flow into early layers and improve training stability. MedNeXt modernizes this design using ConvNeXt-inspired blocks featuring depthwise separable convolutions and inverted bottleneck channel scaling, which increase the effective receptive field while reducing parameter counts. SwinUNETR introduces a hierarchical Swin Transformer encoder that performs local shifted-window multi-head self-attention, capturing long-range dependencies while limiting computational cost; a convolutional U-Net decoder with skip connections reconstructs spatial resolution. MA-SAM adapts the pretrained Segment Anything Model [3] to volumetric medical segmentation by introducing 3D CNN adapters and multi-head attention modules within each Transformer block, while keeping the ViT-Large encoder weights frozen and applying low-rank factorization (FacT) for parameter-efficient fine-tuning. Deep supervision was applied consistently to TractSeg, MedNeXt, and SwinUNETR; MA-SAM was excluded from this scheme as its transformer-based decoder architecture is not readily compatible with intermediate deep supervision.

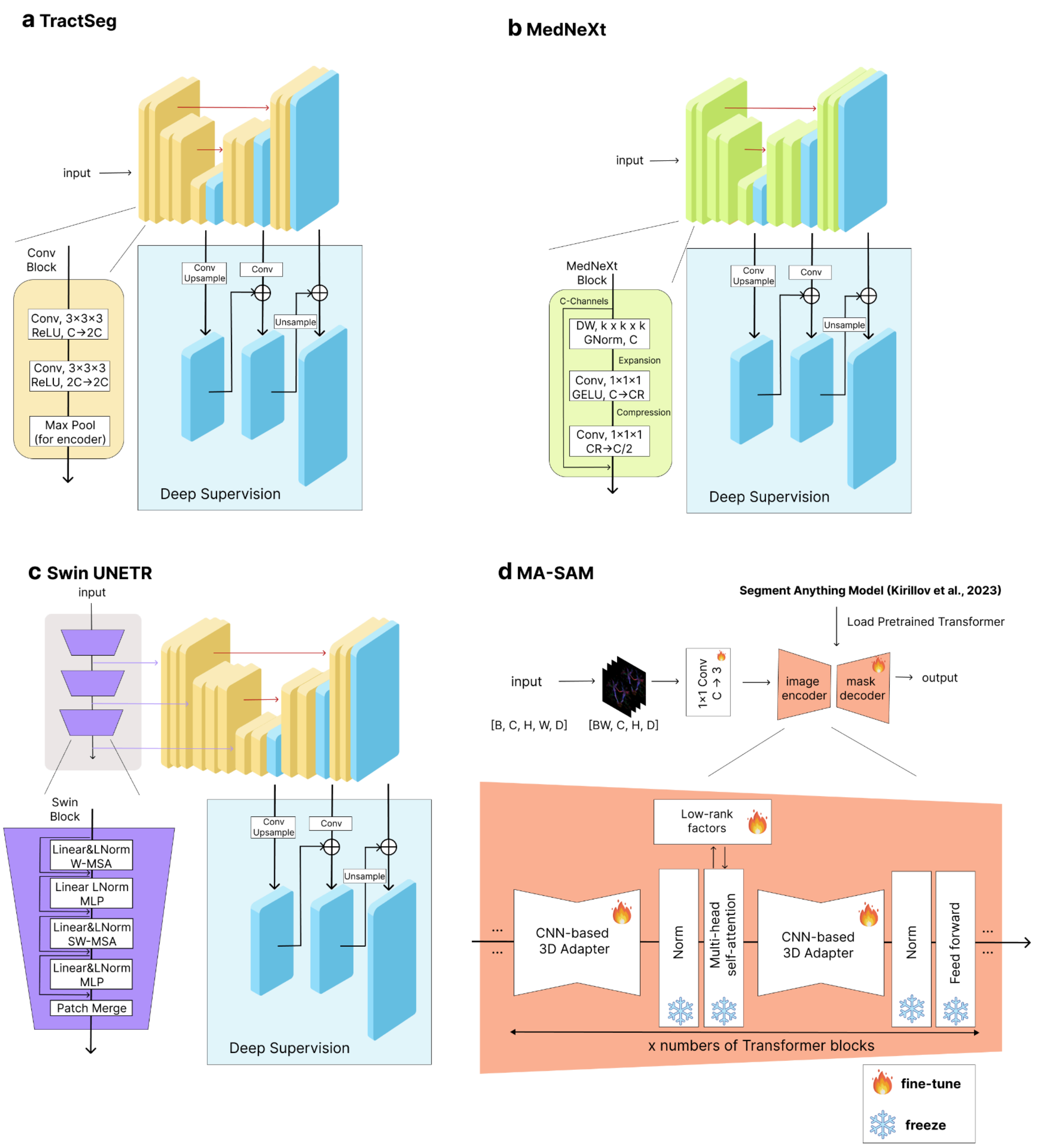


**Supplementary Figure 2. Detailed illustrations of the four deep learning model architectures evaluated for white matter tract segmentation. a.** TractSeg: a 3D U-Net with paired 3×3×3 convolutional encoder blocks, max pooling for downsampling, and skip connections between encoder and decoder stages; intermediate decoder activations (blue blocks) are upsampled and combined with the final output as auxiliary deep supervision losses. **b.** MedNeXt: a ConvNeXt-inspired architecture featuring MedNeXt blocks with depth-wise separable convolutions (DW, k×k×k), grouped normalization (GNorm), GELU activations, and pointwise convolution with inverted bottleneck channel scaling (C→2C). **c.** SwinUNETR: a hybrid architecture with a hierarchical Swin Transformer encoder performing shifted-window multi-head self-attention (W-MSA) with patch merging for downsampling, and a convolutional U-Net decoder with long-range skip connections. **d.** MA-SAM: a ViT-Large encoder initialized from the pretrained Segment Anything Model [3], adapted to 3D volumetric data via CNN-based 3D adapters and multi-head attention modules, with frozen SAM encoder weights fine-tuned through low-rank factorization (FacT). Deep supervision was applied consistently across TractSeg, MedNeXt, and SwinUNETR; MA-SAM was excluded as its transformer-based decoder is not compatible with intermediate supervision.

### Supplementary Result 3: Volumetric context improves TractSeg performance across diverse settings

The original TractSeg was implemented as a 2-dimensional, slice-based model [2]. Here, we developed a 3-dimensional, volume-based variant and tested it across experimental settings (**Supplementary Fig. 3**). Volume-based TractSeg consistently outperformed slice-based TractSeg across most scenarios. Specifically, volume-based TractSeg achieved significantly higher Dice coefficients in 6 of 15 settings (Two-sided t-test, $HCP_{1057}$→$HCP_{1057}$, $p < 0.001$; All Data→$HCP_{1057}$, $p < 0.001$; $PING_{108}$→Cam-$CAN_{288}$, p < 0.001; $HCP_{1057}$→Cam-$CAN_{288}$, $p = 0.026$; $PING_{108}$ →$HCP_{1057}$, $p < 0.001$; Cam-$CAN_{288}$→$HCP_{1057}$ , $p < 0.001$), showed no significant difference in 8 settings, and underperformed slice-based TractSeg in only 1 setting ($HCP_{1057}$→$PING_{108}$; $p < 0.001$). For comparison, we also evaluated SwinUNETR in both slice-based and volume-based versions. Volume-based and slice-based SwinUNETR showed no significant performance difference in 10 of 15 settings. Volume-based SwinUNETR demonstrated superior performance only in two within-domain settings ($HCP_{1057}$→$HCP_{1057}$, $p = 0.016$; All Data→$HCP_{1057}$, $p < 0.001$), whereas slice-based SwinUNETR outperformed its volume-based counterpart in three out-of-domain settings ($HCP_{1057}$&Cam-$CAN_{288}$→$PING_{108}$, $p < 0.001$; Cam-$CAN_{288}$→$HCP_{1057}$, $p < 0.001$); $PING_{108}$&Cam-$CAN_{288}$→$HCP_{1057}$, $p < 0.001$). These results indicate that the performance advantage of volume-based over slice-based processing is architecture-dependent, with TractSeg benefiting more consistently from volumetric context than SwinUNETR.

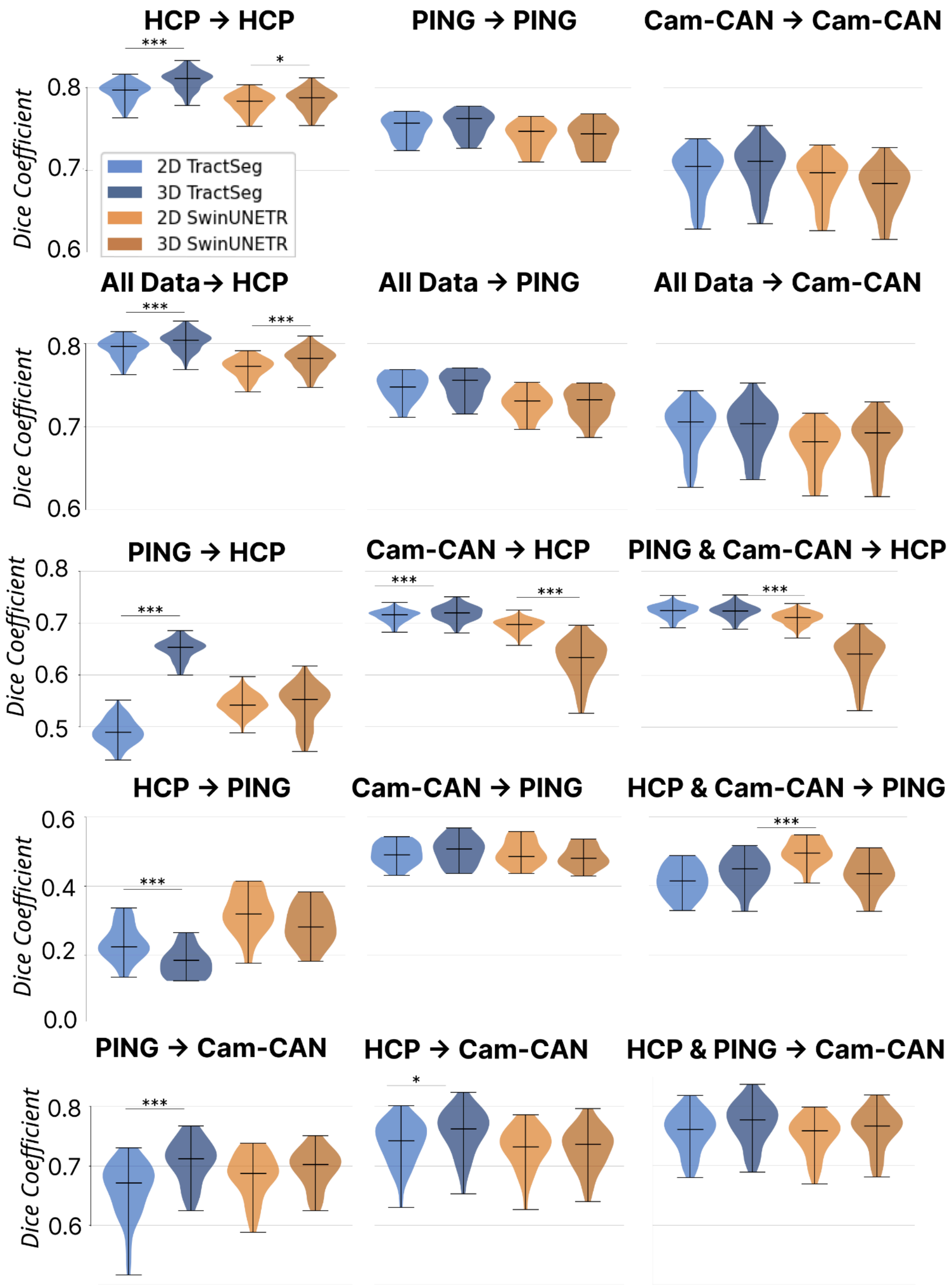


**Supplementary Figure 3. Comparison of segmentation performance between slice-based (2D) and volume-based (3D) implementations of TractSeg and SwinUNETR across all experimental settings.** Violin plots show Dice coefficient distributions for 2D TractSeg (light blue), 3D TractSeg (dark blue), 2D SwinUNETR (light orange), and 3D SwinUNETR (dark orange) in each training–testing scenario. Panels are organized by test dataset and grouped into within-domain settings (top two rows: HCP→HCP, PING→PING, Cam-CAN→Cam-CAN, All Data→HCP, All Data→PING, All Data→Cam-CAN) and out-of-domain generalization settings (bottom three rows). Each panel title denotes the training dataset (left of the arrow) and test dataset (right of the arrow). Asterisks indicate statistically significant differences between 2D and 3D variants of the same architecture (two-sided t-test; $^*p < 0.05$, $^{***}p < 0.001$). Horizontal lines within violins indicate median values; error bars represent the interquartile range.

**Supplementary Result 4: Tract-specific segmentation performance across all benchmark experiments**

Segmentation accuracy varied substantially across the 61 white matter tracts in all experimental conditions, revealing consistent patterns of tract-specific difficulty that cut across model architectures and datasets (**Supplementary Fig. 4**). Large, well-defined projection tracts, such as the corticospinal tract and corpus callosum segments, consistently achieved the highest Dice coefficients (>0.80 in within-domain HCP settings), while smaller association tracts with greater anatomical variability, such as the uncinate fasciculus and posterior arcuate, exhibited lower and more variable performance. Tracts present exclusively in the Brainlife WMA parcellation, including cerebellar tracts, optic radiation subdivisions (Meyer's and Baum's loops), and posterior association tracts (pArc, VOF, TPC), generally showed lower performance than tracts shared with the TractSeg labeling scheme, reflecting their finer anatomical specificity and limited representation in training data (see **Supplementary Table 1** for one-to-one correspondence of two different labeling schemes). These tract-level disparities were broadly consistent across models and settings, indicating that intrinsic tract properties, primarily volume and anatomical distinctiveness, are the primary drivers of segmentation difficulty, rather than model architecture or training condition per se (see also **Supplementary Fig. 5**).

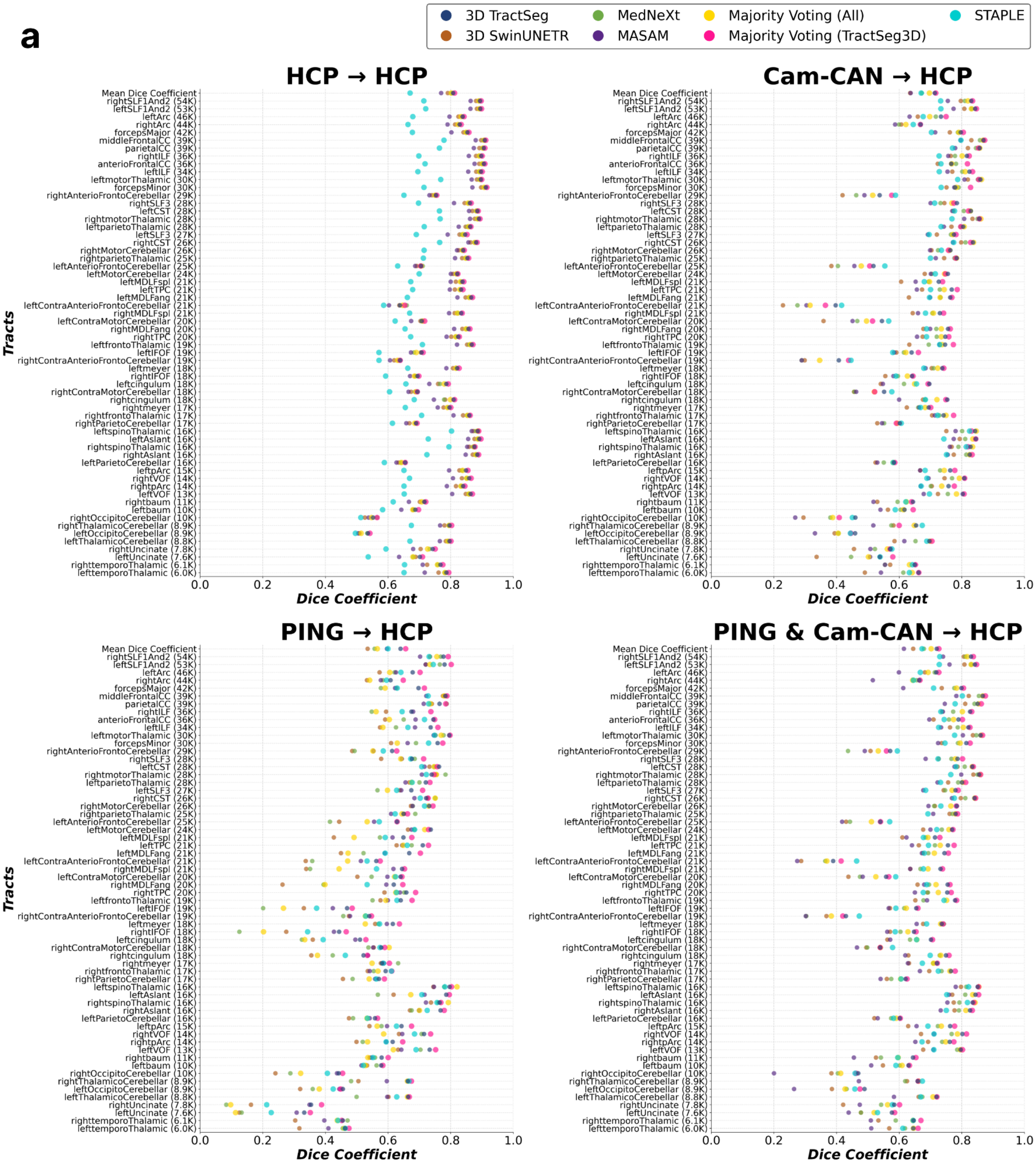
a
3D TractSeg
3D SwinUNETR
MedNeXt
MASAM
Majority Voting (All)
Majority Voting (TractSeg3D)
STAPLE
HCP → HCP
Cam-CAN → HCP
PING → HCP
PING & Cam-CAN → HCP
Tracts
Dice Coefficient
Mean Dice Coefficient
rightSLF1And2 (54K)
leftSLF1And2 (53K)
leftArc (46K)
rightArc (44K)
forcepsMajor (42K)
middleFrontalCC (39K)
parietalCC (39K)
rightILF (36K)
anterioFrontalCC (36K)
leftILF (34K)
leftmotorThalamic (30K)
forcepsMinor (30K)
rightAnterioFrontoCerebellar (29K)
rightSLF3 (28K)
leftCST (28K)
rightmotorThalamic (28K)
leftparietoThalamic (28K)
leftSLF3 (27K)
rightCST (26K)
rightMotorCerebellar (26K)
rightparietoThalamic (25K)
leftAnterioFrontoCerebellar (25K)
leftMotorCerebellar (24K)
leftMDLFspl (21K)
leftTPC (21K)
leftMDLFang (21K)
leftContraAnterioFrontoCerebellar (21K)
rightMDLFspl (21K)
leftContraMotorCerebellar (20K)
rightMDLFang (20K)
rightTPC (20K)
leftfrontoThalamic (19K)
leftIFOF (19K)
rightContraAnterioFrontoCerebellar (19K)
leftmeyer (18K)
rightIFOF (18K)
leftcingulum (18K)
rightContraMotorCerebellar (18K)
rightcingulum (18K)
rightmeyer (17K)
rightfrontoThalamic (17K)
rightParietoCerebellar (17K)
leftspinoThalamic (16K)
leftAslant (16K)
rightspinoThalamic (16K)
rightAslant (16K)
leftParietoCerebellar (16K)
leftpArc (15K)
rightVOF (14K)
rightpArc (14K)
leftVOF (13K)
rightbaum (11K)
leftbaum (10K)
rightOccipitoCerebellar (10K)
rightThalamicoCerebellar (8.9K)
leftOccipitoCerebellar (8.9K)
leftThalamicoCerebellar (8.8K)
rightUncinate (7.8K)
leftUncinate (7.6K)
righttemporoThalamic (6.1K)
lefttemporoThalamic (6.0K)
0.0
0.2
0.4
0.6
0.8
1.0

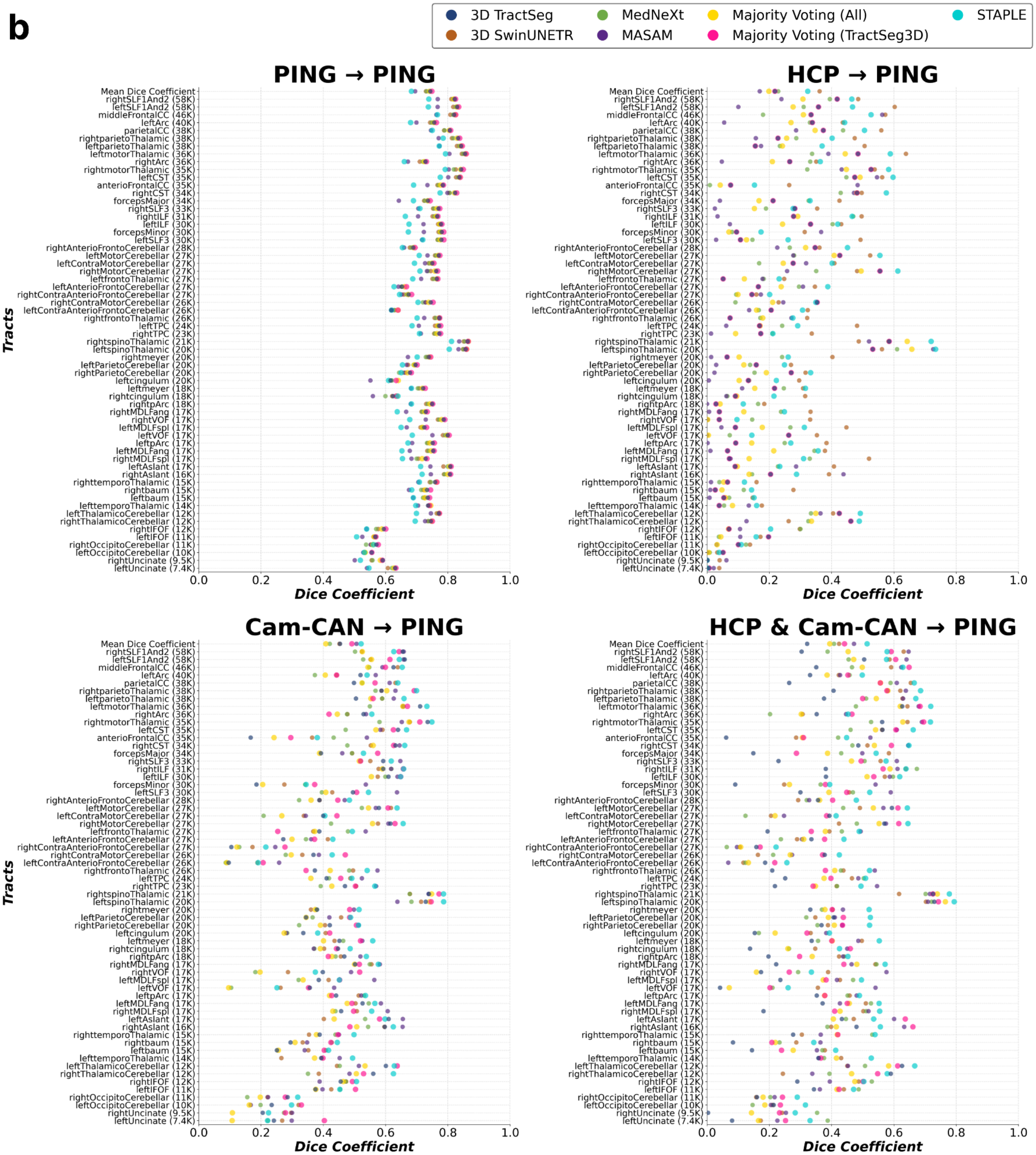
b
3D TractSeg
3D SwinUNETR
MedNeXt
MASAM
Majority Voting (All)
Majority Voting (TractSeg3D)
STAPLE
PING → PING
HCP → PING
Cam-CAN → PING
HCP & Cam-CAN → PING
Tracts
Dice Coefficient
0.0 0.2 0.4 0.6 0.8 1.0
Mean Dice Coefficient
rightSLF1And2 (58K)
leftSLF1And2 (58K)
middleFrontalCC (46K)
leftArc (40K)
parietalCC (38K)
rightparietoThalamic (38K)
leftparietoThalamic (38K)
leftmotorThalamic (36K)
rightArc (36K)
rightmotorThalamic (35K)
leftCST (35K)
anterioFrontalCC (35K)
rightCST (34K)
forcepsMajor (34K)
rightSLF3 (33K)
rightILF (31K)
leftILF (30K)
forcepsMinor (30K)
leftSLF3 (30K)
rightAnterioFrontoCerebellar (28K)
leftMotorCerebellar (27K)
leftContraMotorCerebellar (27K)
rightMotorCerebellar (27K)
leftfrontoThalamic (27K)
leftAnterioFrontoCerebellar (27K)
rightContraAnterioFrontoCerebellar (27K)
rightContraMotorCerebellar (26K)
leftContraAnterioFrontoCerebellar (26K)
rightfrontoThalamic (26K)
leftTPC (24K)
rightTPC (23K)
rightspinoThalamic (21K)
leftspinoThalamic (20K)
rightmeyer (20K)
leftParietoCerebellar (20K)
rightParietoCerebellar (20K)
leftcingulum (20K)
leftmeyer (18K)
rightcingulum (18K)
rightpArc (18K)
rightMDLFang (17K)
rightVOF (17K)
leftMDLFspl (17K)
leftVOF (17K)
leftpArc (17K)
leftMDLFang (17K)
rightMDLFspl (17K)
leftAslant (17K)
rightAslant (16K)
righttemporoThalamic (15K)
rightbaum (15K)
leftbaum (15K)
lefttemporoThalamic (14K)
leftThalamicoCerebellar (12K)
rightThalamicoCerebellar (12K)
rightIFOF (12K)
leftIFOF (11K)
rightOccipitoCerebellar (11K)
leftOccipitoCerebellar (10K)
rightUncinate (9.5K)
leftUncinate (7.4K)

c

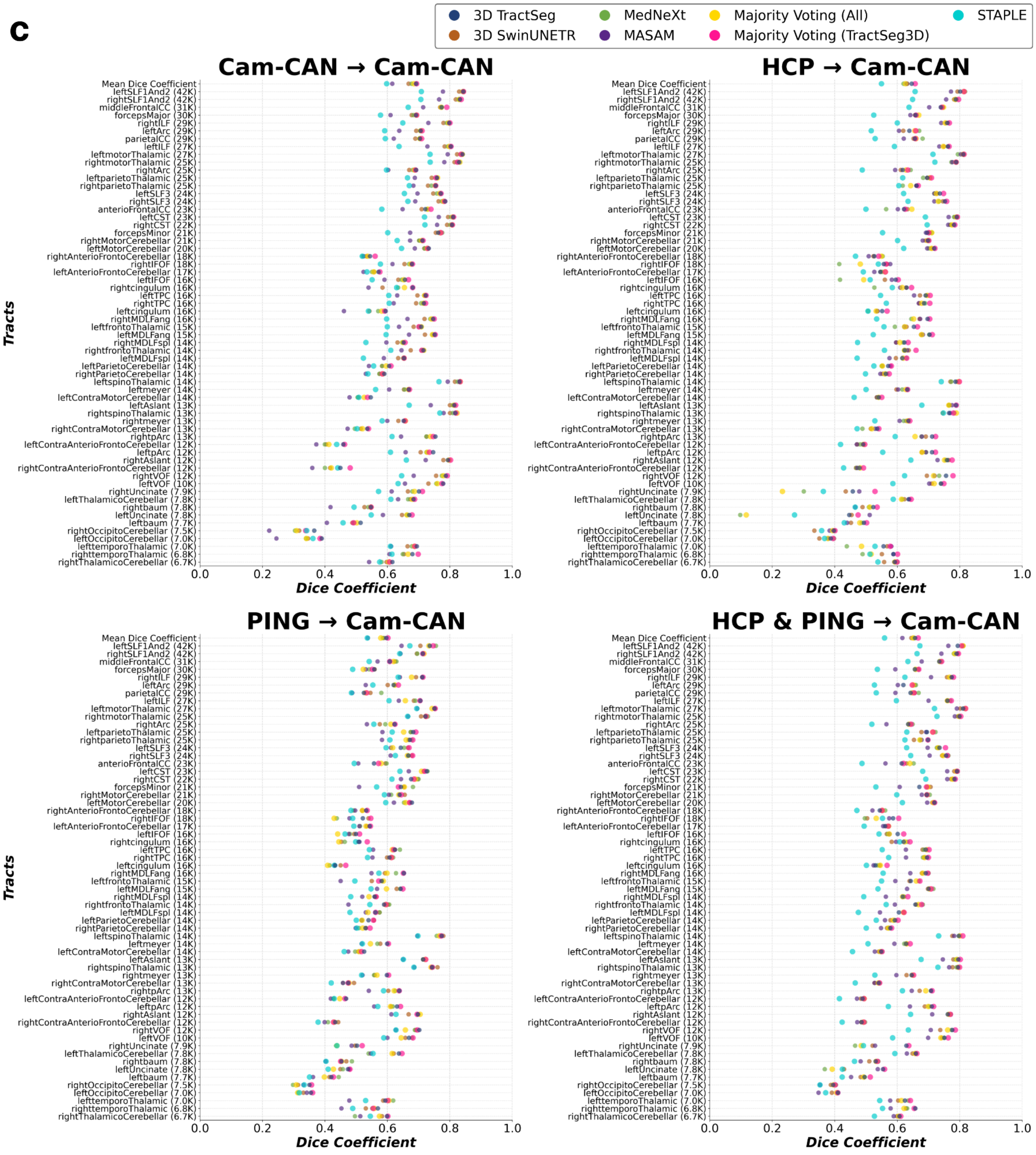


**Supplementary Figure 4. Tract-specific segmentation performance across 12 experimental settings.** Dice coefficients for each of the 61 white matter tracts are shown for all benchmark models (3D TractSeg, MedNeXt, 3D SwinUNETR, and MASAM) and ensemble-based approaches (Majority Voting-based approaches and STAPLE). Tracts are ordered along the y-axis by descending average voxel count, computed by averaging ground-truth tract masks across all subjects within each dataset and thresholding the resulting mean image at 0.5. **a.** Settings in which HCP serves as the test set. **b.** Settings in which PING serves as the test set. **c.** Settings in which Cam-CAN serves as the test set.

#### Supplementary Result 5: Tracts with lower volume have lower prediction accuracy.

Segmentation accuracy varied substantially across the 61 white matter tracts (**Supplementary Fig. 4**). To assess whether tract volume underlies this variability, we correlated voxel count with Dice coefficient for 3D TractSeg across three within-domain settings (**Supplementary Fig. 5**). Larger tracts consistently achieved higher Dice scores in all three datasets, with moderate positive correlations: r = 0.50 ($p = 3.96 \times 10^{-5}$) for $HCP_{1057} \rightarrow HCP_{1057}$, r = 0.52 ($p = 1.54 \times 10^{-5}$) for $Cam\text{-}CAN_{288} \rightarrow Cam\text{-}CAN_{288}$, and r = 0.56 ($p = 3.07 \times 10^{-6}$) for $PING_{108} \rightarrow PING_{108}$.

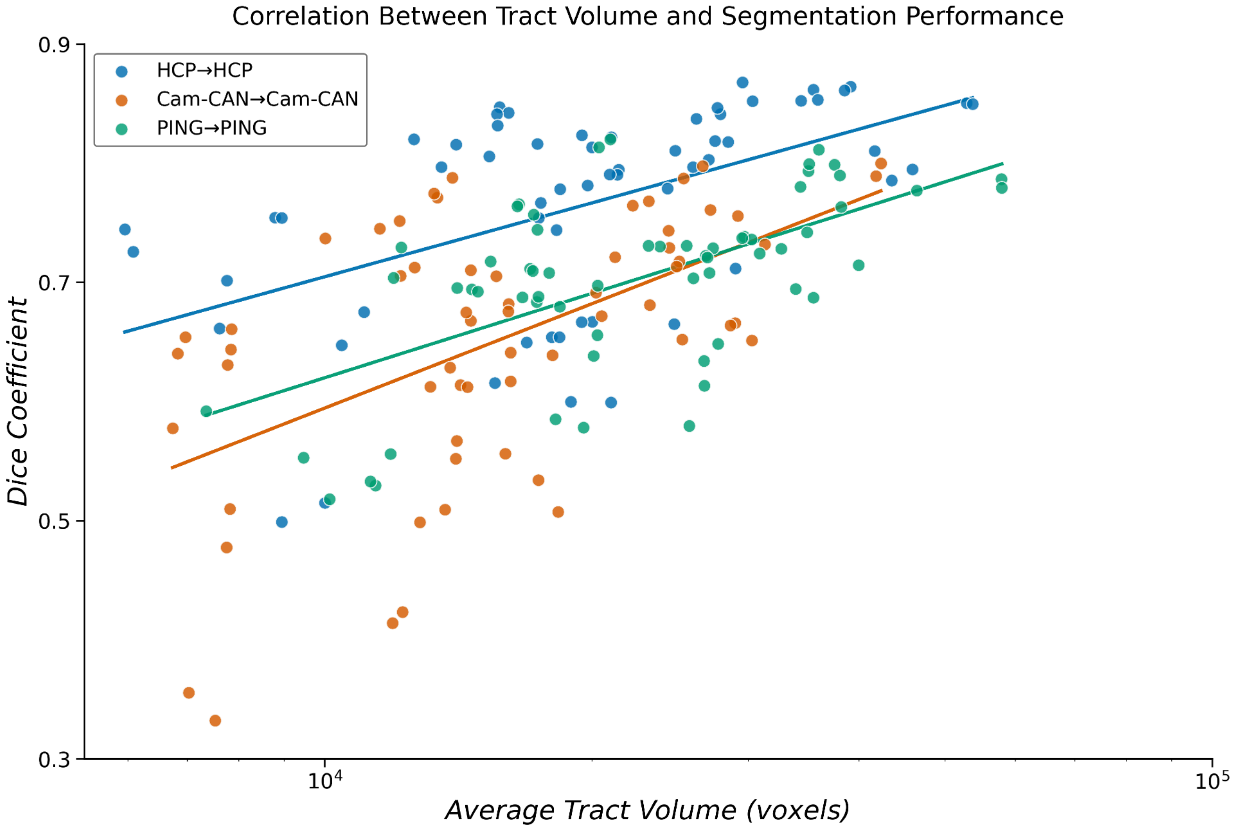


**Supplementary Figure 5.** Relationship between tract volume and tract-wise segmentation performance across within-domain evaluations. Each point represents one of the 61 tracts, with the x-axis showing the average tract volume in voxels on a logarithmic scale and the y-axis showing the tract-wise Dice coefficient (3D TractSeg). Results are shown for three within-domain settings (HCP→HCP, Cam-CAN→Cam-CAN, and PING→PING). Average tract volumes were counted from the reference bundle masks of each dataset, and Dice scores were obtained from the corresponding tract-wise test results. Solid lines indicate linear fits estimated in log-volume space to visualize the trend between tract size and segmentation accuracy.

### Supplementary Result 6: Comparing model performance between a small-curated dataset and a natural uncurated dataset

In **Supplementary Fig.6**, we compared the model using two different sets of tract masks: 72 tract masks for $HCP_{105}$ [2] and our 61 tract masks [4,5]. For a fair comparison between the distinct tracts, we used the same subjects from the 105 HCP dataset, naming the processed data $HCP_{Wasserthal}$(105) and our approach $HCP_{Ours}$(105). Since the authors did not offer the DWI data, we processed it using the DWI pipeline described in the previous section. We named the case where we processed the entire dataset of 1057 subjects as $HCP_{Ours}$(1057). The data split adhered to the original protocol, comprising 63 subjects for training, 21 for validation, and 21 for testing, with cross-validation performed across multiple random splits.

Models trained on $HCP_{Wasserthal}$(105) exhibited the same performance ranking observed in our homogeneous training conditions: TractSeg, MedNeXt, Swin-UNETR, and MA-SAM, in descending order. Overall performance on $HCP_{Wasserthal}$(105) ranged from 0.81 to 0.84 Dice coefficient across models, while our $HCP_{Ours}$(105) dataset yielded slightly lower performance, ranging from 0.75 to 0.79. (Note that direct comparison of these absolute values should be interpreted cautiously, as the two datasets differ in the number and specific definitions of tract labels.) Notably, expanding the training set from 105 to 1,057 subjects using our automated pipeline ($HCP_{Ours}$(1057) resulted in consistent performance improvements across all models compared to $HCP_{Ours}$(105). This improvement demonstrates that even within a single homogeneous dataset, increasing sample size enhances model performance across all architectures, suggesting that the benefits of larger training sets extend beyond capturing population diversity to improving statistical robustness of learned representations.

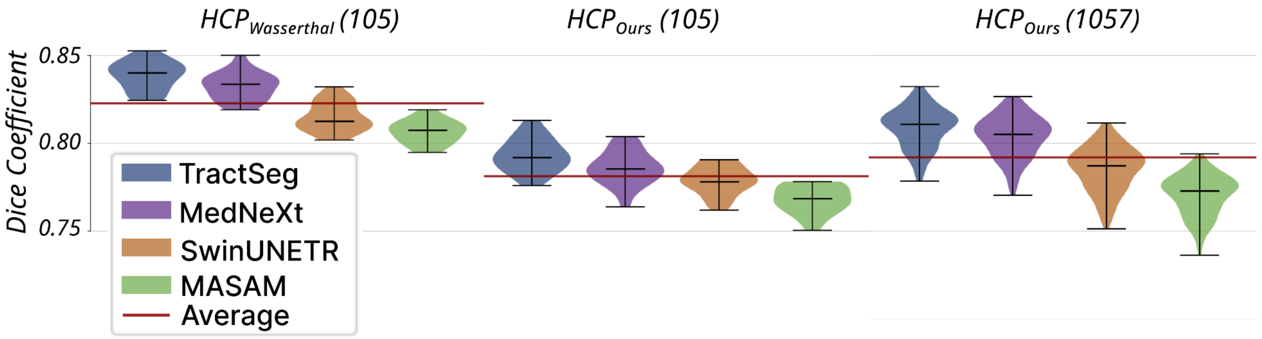


**Supplementary Figure 6. Comparison of model segmentation performance between a curated benchmark dataset by ref 1 and by our automated pipeline.** Violin plots show Dice coefficient distributions for TractSeg (blue), MedNeXt (purple), SwinUNETR (orange), and MA-SAM (green) across three conditions: $HCP_{Wasserthal}$(105), which uses the 72 tract labels from the original TractSeg study [2] processed with the published pipeline; $HCP_{Ours}$(105), which applies our automated brainlife.io pipeline to the same 105 subjects with 61 tract labels; and $HCP_{Ours}$(1057), which scales up our automated pipeline to the full 1,057-subject HCP cohort. The red horizontal line indicates the mean Dice coefficient across models for each condition. All experiments used the same data split as the protocol in ref. 1 (63 subjects for training, 21 for validation, 21 for testing), with cross-validation performed across multiple random splits. Note that direct comparison of absolute Dice values between $HCP_{Wasserthal}$(105) and $HCP_{Ours}$(105) should be interpreted with caution, as the two datasets differ in the number and anatomical definitions of tract labels.

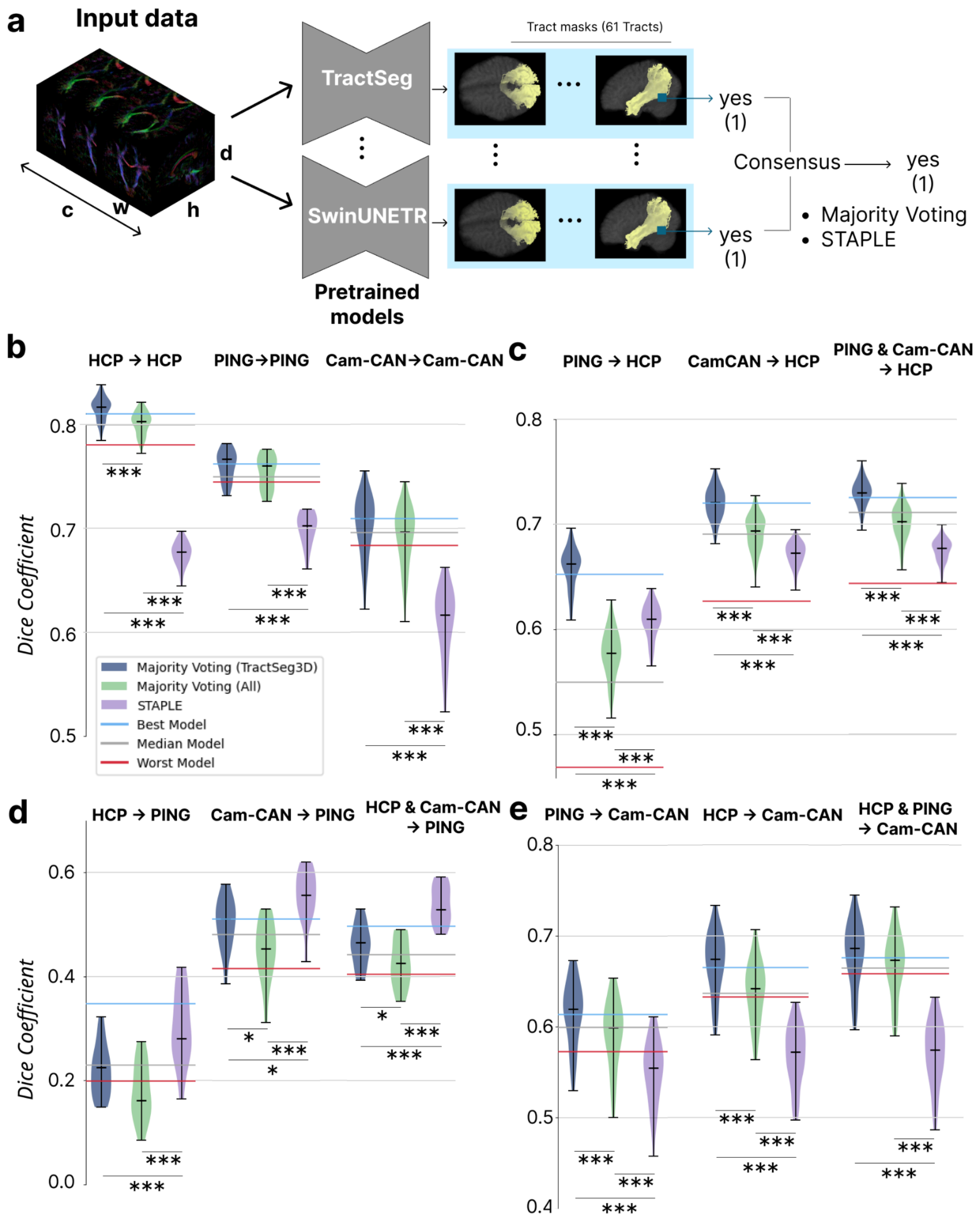


**Supplementary Figure 7. Ensemble methods improve robustness across generalization settings. a.** Input diffusion MRI data (left) are processed through pretrained segmentation models, each predicting tract masks for 61 WMT (middle). Consensus tract masks are generated via ensemble integration using either majority voting or the STAPLE (Simultaneous Truth and Performance Level Estimation) algorithm (right). **b.** Within-domain ensemble performance across three single-dataset training settings. Violin plots compare STAPLE integration across five models (purple), majority voting across five models (dark blue), and majority voting across three TractSeg3D models with different data splits (green). Horizontal lines indicate best (blue), median (grey), and worst (red) individual model performance. **c–e.** Out-of-domain generalization of ensemble models. Dice coefficients are shown for transfers to $HCP_{1057}$ (c), Cam-$CAN_{288}$ (d), and $PING_{108}$ (e) from various source datasets. Statistical significance: *$p < 0.05$, **$p < 0.01$,***$p < 0.001$ (Pair-wise t-test with Bonferroni correction). Error bars represent interquartile range; horizontal lines within violins indicate median values.

### Supplementary Result 7: Ensemble methods for improved generalization

Our previous analyses demonstrated that model performance can vary across data settings; no single model architecture was consistently superior to others. In within-domain evaluations, CNN-based models, particularly TractSeg and MedNeXt, outperformed transformer-based approaches. In out-of-domain transfers, however, performance depended on the target dataset; transformers showed advantages in more challenging shifts, such as SwinUNETR in $HCP_{1057}$ to $PING_{108}$, whereas CNNs performed better in others, such as TractSeg in Cam-$CAN_{288}$ to $HCP_{1057}$. This architecture-specific sensitivity to distributional shifts indicates that no single model is optimal across deployment scenarios.

We tested whether ensemble methods could leverage the new benchmark dataset to achieve more robust generalization to unseen datasets (**Supplementary Fig. 7a**). We implemented two ensemble strategies: majority voting [6] and STAPLE [7]. Majority voting generates a consensus segmentation by assigning each voxel the label predicted by most models, whereas STAPLE produces a probabilistic consensus by jointly estimating the true segmentation and the sensitivity and specificity of each model, thereby weighting contributions according to inferred model performance. We applied these two ensemble methods across five models: slice-based TractSeg, volume-based TractSeg, slice-based SwinUNETR, volume-based SwinUNETR, and volume-based MedNeXt (MA-SAM was excluded due to consistently poor performance in all previous experiments). Additionally, using the overall best-performing model (TractSeg 3D), we generated ensemble predictions on a fixed test set by applying majority voting across three models trained on different splits of the non-test subjects, thereby evaluating the robustness of performance to variability in the training data.

*Within-Domain Settings* (**Supplementary Fig. 7b**): Compared to individual models' performance, majority voting performed consistently between the median and the best individual model. Furthermore, the TractSeg3D ensemble matched or slightly exceeded the best individual model (i.e., TractSeg3D). STAPLE underperformed across conditions, in some cases falling below the weakest individual model.

*Out-of-Domain Settings* (**Supplementary Fig. 7c–e**): We next examined how data distributional shifts affect ensemble results. Ensemble behavior diverged between transfer targets. When transferring to $HCP_{1057}$ (**Supplementary Fig. 7c**), the ensemble ranking depended on the source dataset. In the $PING_{108} \rightarrow HCP_{1057}$ setting, STAPLE significantly outperformed majority voting (STAPLE Dice coefficient 0.61 versus majority voting 0.58; $p < 0.001$), one of the settings in which transformer-based models showed relatively higher performance. In the remaining $HCP_{1057}$ transfers (Cam-$CAN_{288} \rightarrow HCP_{1057}$ and $PING_{108}$&Cam-$CAN_{288} \rightarrow HCP_{1057}$), however, majority voting regained superiority over STAPLE. When transferring to $PING_{108}$ (**Supplementary Fig. 7d**), STAPLE consistently outperformed majority voting, particularly in the most challenging $HCP_{1057} \rightarrow PING_{108}$ scenario, where individual models exhibited severe performance degradation (STAPLE Dice coefficient 0.30 versus majority voting 0.18, $p < 0.001$). The Cam-$CAN_{288} \rightarrow PING_{108}$ and $HCP_{1057}$&Cam-$CAN_{288} \rightarrow PING_{108}$ transfers exhibited a similar but attenuated pattern (Cam-$CAN_{288} \rightarrow PING_{108}$: STAPLE 0.56 versus majority voting 0.45, $p < 0.001$; $HCP_{1057}$&Cam-$CAN_{288} \rightarrow PING_{108}$: STAPLE 0.54 versus majority voting 0.42, $p < 0.001$). By contrast, when transferring to Cam-$CAN_{288}$ (**Supplementary Fig. 7e**), majority voting consistently performed between the median and the best individual models across all source datasets ($HCP_{1057}$, $PING_{108}$, $HCP_{1057}$&$PING_{108}$), whereas STAPLE underperformed relative to the worst individual model, suggesting that Cam-$CAN_{288}$'s moderate difficulty does not necessitate weighted integration. Collectively, these results suggest that STAPLE achieves greater performance gains over simple averaging in challenging scenarios where transformer-based architectures previously demonstrated advantages over CNN-based models.

### Supplementary Result 8: Ensemble performances depending on thresholds

To investigate why STAPLE performs worse than individual models in within-domain settings, yet outperforms majority voting in out-of-domain settings, we examined how performance varies with the final threshold. The prediction maps from both STAPLE and majority voting are probability values ranging from 0 to 1, and all prior analyses were conducted using a threshold of 0.5. The analysis revealed that STAPLE was largely insensitive to threshold changes, whereas majority voting showed substantial performance variation depending on the threshold (**Supplementary Fig. 8**). In settings where majority voting previously outperformed STAPLE, majority voting tended to peak at a threshold of 0.5 while STAPLE maintained relatively stable performance regardless of the threshold (11 out of 15 settings). In contrast, in settings where STAPLE previously outperformed majority voting (other datasets to PING inference, and PING to HCP), both methods achieved optimal performance at very low thresholds (around 0.1); however, majority voting experienced a much steeper performance decline than STAPLE as the threshold increased, resulting in STAPLE outperforming majority voting at the 0.5 threshold.

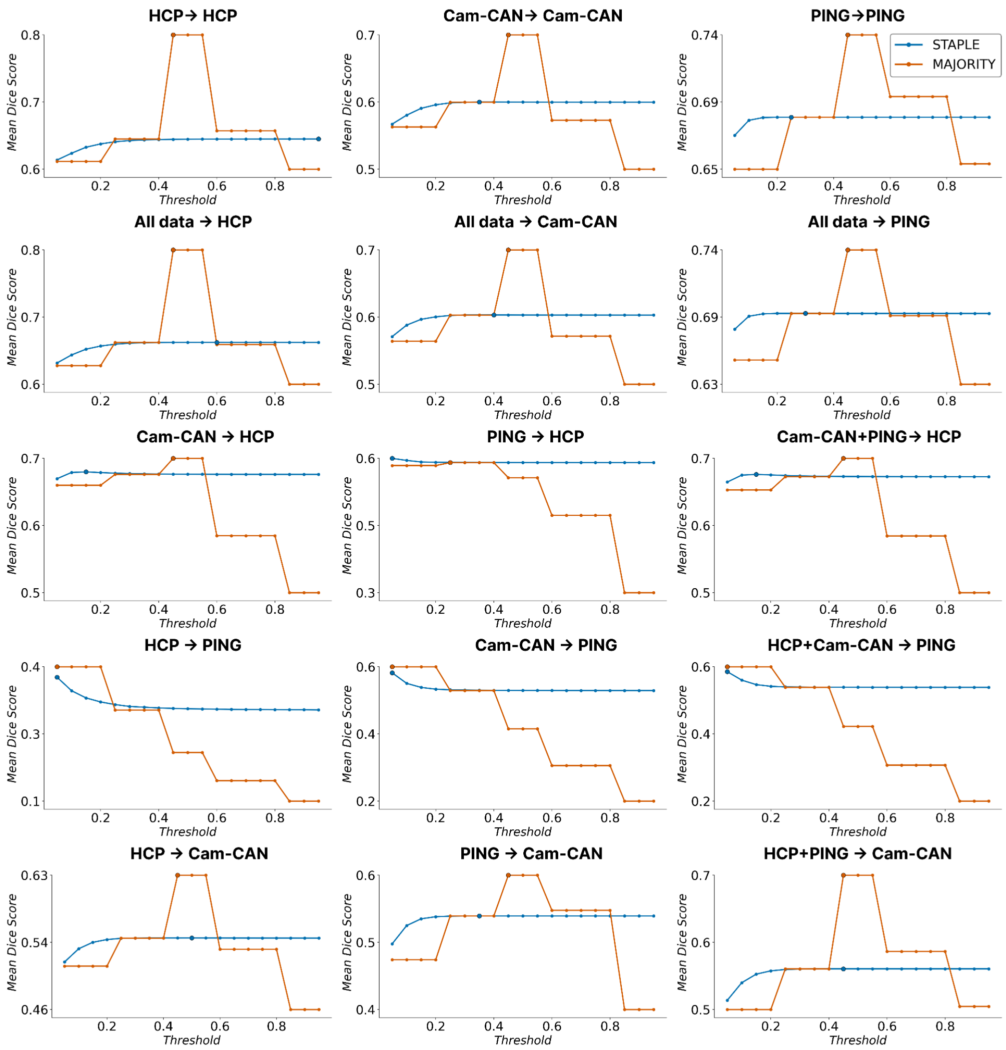


**Supplementary Figure 8. Performance of STAPLE and majority voting ensemble methods across different probability thresholds in within-domain and out-of-domain settings.** Dice similarity coefficient (DSC) is plotted as a function of the binarization threshold (0 to 1) for STAPLE and majority voting across all evaluated settings. In within-domain settings where majority voting outperformed STAPLE, it consistently peaked near 0.5, whereas STAPLE exhibited relatively stable performance across thresholds (11 out of 15 settings). In out-of-domain settings where STAPLE outperformed majority voting (other datasets to PING inference, and PING to HCP), both methods achieved optimal performance at substantially lower thresholds (around 0.1); however, majority voting showed a markedly steeper decline in performance as the threshold increased, whereas STAPLE remained comparatively stable, resulting in higher DSC for STAPLE at the standard 0.5 threshold.

### Supplementary Result 9. Qualitative evaluation shows the prediction quality depending on the tract

To provide deeper insights into model performance beyond quantitative metrics, we conducted a qualitative analysis of predicted tract masks across representative within-domain and out-of-domain scenarios. **Supplementary Fig. 9** presents segmentation results for two white matter tracts with contrasting characteristics: the right corticospinal tract (CST), a large, well-defined projection tract that demonstrated high performance across all methods (mean Dice >0.80), and the left uncinate fasciculus, a smaller association tract with more anatomical variability that showed relatively lower performance in our quantitative evaluation (mean Dice ~0.65). In within-domain settings (left columns), all methods produced segmentations that closely matched the reference masks for both tracts. For the CST, both majority voting across TractSeg models and across all five models captured the characteristic superior-inferior trajectory from the motor cortex to the brainstem, with minimal differences. STAPLE appeared very similar to the other two approaches, but it overestimated tract size compared to the other approaches and thus showed lower performance in the Dice coefficient compared to the other approaches (**Supplementary Fig. 9**). The uncinate fasciculus, despite its smaller size and curved anatomy, was similarly well-reconstructed across all ensemble methods in within-domain conditions. These results visually confirm our quantitative findings that ensemble methods perform comparably to the best individual models when trained and tested on matched populations.

In contrast, out-of-domain settings revealed substantial performance degradation and notable differences between ensemble strategies. When transferring from $PING_{108}$ to $HCP_{1057}$ (top right), the CST remained relatively well-preserved across all methods, though with slightly reduced coverage in the superior portions. However, for the uncinate fasciculus, the performance gap between methods became pronounced: majority voting from TractSeg models and all five models produced severely fragmented masks with substantial missing anatomy, while STAPLE maintained more complete tract coverage, albeit still reduced compared to within-domain predictions.

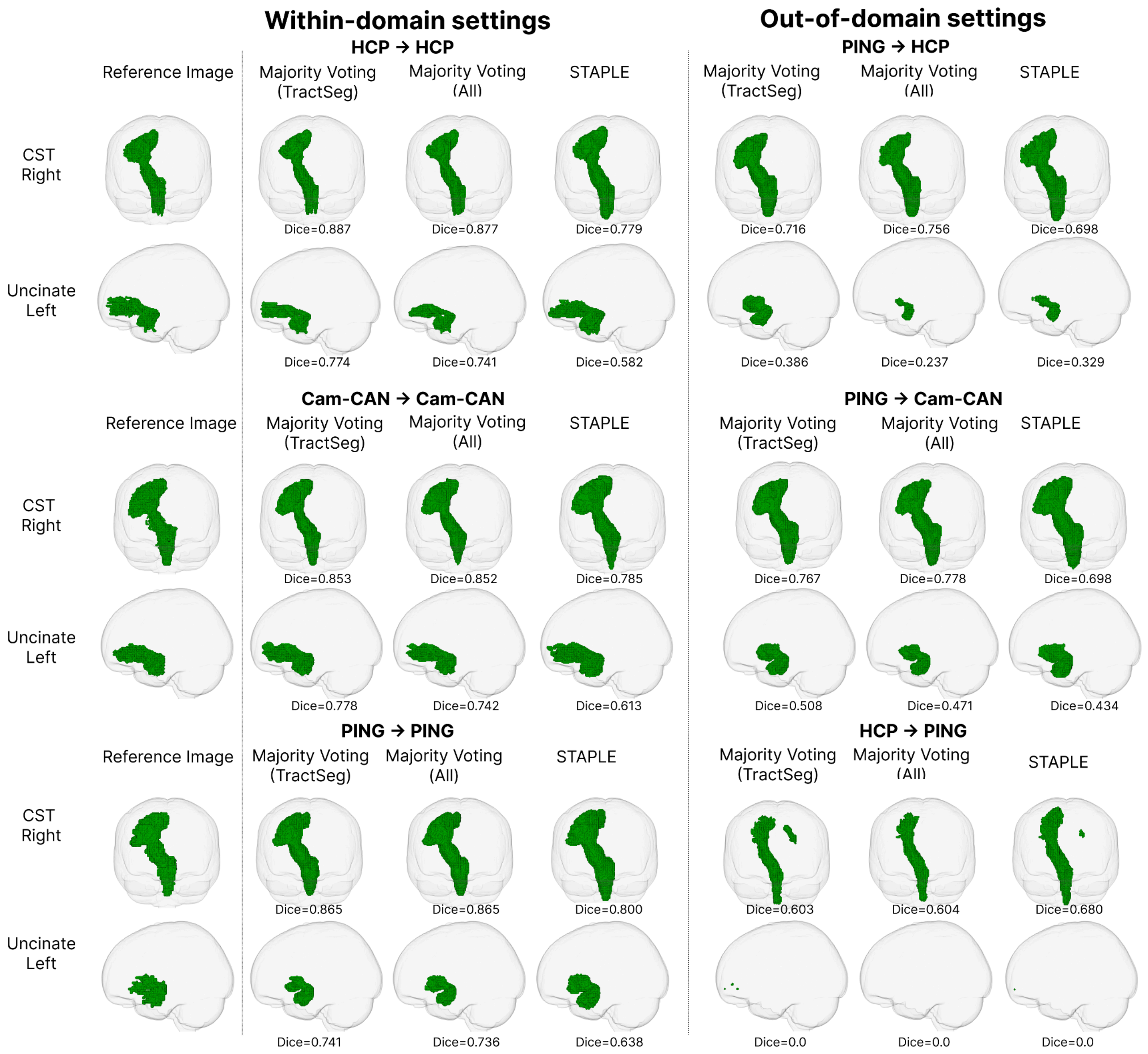


**Supplementary Figure 9. Qualitative Evaluation of each modeling approach across different settings, models, and tracts.** Representative segmentation results for two white matter tracts with distinct anatomical characteristics across six experimental scenarios. A threshold of 0.5 was used for ensemble predictions. Left columns show within-domain settings where training and test data are from the same dataset; right columns show out-of-domain settings where models are tested on populations different from training. Top row: right corticospinal tract (CST), a large projection tract. Bottom row: left uncinate fasciculus, a smaller association tract with greater anatomical variability. For each scenario, segmentations are shown for the reference (ground truth) mask, majority voting from three TractSeg3D models that differ only in data splits, majority voting from all five models (TractSeg3D volume and slice, Swin-UNETR volume and slice, excluding MA-SAM), and a STAPLE-weighted ensemble.

### Supplementary Tables

| Supplementary Table 1. Correspondence between Brainlife WMA and TractSeg tracts. | | | |
|---|---|---|---|
| | **brainlife.io** | **TracSeg** | **Tract description** |
| **1** | anterioFrontalCC | CC_1\|CC_2 | anterior frontal segment of the corpus callosum |
| **2** | forcepsMajor | CC_7 | forceps major (posterior callosal fibers) |
| **3** | forcepsMinor | CC_1\|CC_2\|frontal WM fan-out | forceps minor (anterior callosal fibers) |
| **4** | leftAnterioFrontoCerebellar | only brainlife | left hemisphere anterior fronto-cerebellar tract |
| **5** | leftArc | AF_left | left hemisphere arcuate fasciculus |
| **6** | leftAslant | only brainlife | left hemisphere frontal aslant tract |
| **7** | leftCST | CST_left | left hemisphere corticospinal tract |
| **8** | leftContraAnterioFrontoCerebellar | only brainlife | left hemisphere contralateral anterior fronto-cerebellar tract |
| **9** | leftContraMotorCerebellar | only brainlife | left hemisphere contralateral motor cerebellar tract |
| **10** | leftIFOF | IFO_left | left hemisphere inferior fronto-occipital fasciculus |
| **11** | leftILF | ILF_left | left hemisphere inferior longitudinal fasciculus |
| **12** | leftMDLFang | only brainlife | left hemisphere middle longitudinal fasciculus, angular-gyrus branch |
| **13** | leftMDLFspl | only brainlife | left hemisphere middle longitudinal fasciculus, superior-parietal-lobule branch |
| **14** | leftMotorCerebellar | only brainlife | left hemisphere motor cerebellar tract |
| **15** | leftOccipitoCerebellar | only brainlife | left hemisphere occipito-cerebellar tract |
| **16** | leftParietoCerebellar | only brainlife | left hemisphere parieto-cerebellar tract |
| **17** | leftSLF1And2 | SLF_I_left\|SLF_II_left | left hemisphere superior longitudinal fasciculus I and II (combined) |
| **18** | leftSLF3 | SLF_III_left | left hemisphere superior longitudinal fasciculus III |
| **19** | leftTPC | only brainlife | left hemisphere temporo-parietal connection |
| **20** | leftThalamicoCerebellar | only brainlife | left hemisphere thalamo-cerebellar tract |
| **21** | leftUncinate | UF_left | left hemisphere uncinate fasciculus |
| **22** | leftVOF | only brainlife | left hemisphere vertical occipital fasciculus |
| **23** | leftbaum | only brainlife | left hemisphere Baum's loop (dorsal optic radiation) |
| **24** | leftcingulum | CG_left | left hemisphere cingulum bundle |
| **25** | leftfrontoThalamic | only brainlife | left hemisphere fronto-thalamic tract |
| **26** | leftmeyer | only brainlife | left hemisphere Meyer's loop (ventral optic radiation) |
| **27** | leftmotorThalamic | only brainlife | left hemisphere motor-thalamic tract |
| **28** | leftpArc | only brainlife | left hemisphere posterior arcuate fasciculus |
| **29** | leftparietoThalamic | only brainlife | left hemisphere parieto-thalamic tract |
| **30** | leftspinoThalamic | only brainlife | left hemisphere spinothalamic tract |
| **31** | lefttemporoThalamic | only brainlife | left hemisphere temporo-thalamic tract |
| **32** | middleFrontalCC | CC_3\|CC_4 | middle frontal segment of corpus callosum |
| **33** | parietalCC | CC_6 | parietal segment of corpus callosum |
| **34** | rightAnterioFrontoCerebellar | only brainlife | right hemisphere anterior fronto-cerebellar tract |
| **35** | rightArc | AF_right | right hemisphere arcuate fasciculus |

| **Supplementary Table 1.** Correspondence between Brainlife WMA and TractSeg tracts. | | | |
|---|---|---|---|
| **36** | rightAslant | only brainlife | right hemisphere frontal aslant tract |
| **37** | rightCST | CST_right | right hemisphere corticospinal tract |
| **38** | rightContraAnterioFrontoCerebellar | only brainlife | right hemisphere contralateral anterior fronto-cerebellar tract |
| **39** | rightContraMotorCerebellar | only brainlife | right hemisphere contralateral motor cerebellar tract |
| **40** | rightIFOF | IFO_right | right hemisphere inferior fronto-occipital fasciculus |
| **41** | rightILF | ILF_right | right hemisphere inferior longitudinal fasciculus |
| **42** | rightMDLFang | only brainlife | right hemisphere middle longitudinal fasciculus, angular-gyrus branch |
| **43** | rightMDLFspl | only brainlife | right hemisphere middle longitudinal fasciculus, superior-parietal-lobule branch |
| **44** | rightMotorCerebellar | only brainlife | right hemisphere motor cerebellar tract |
| **45** | rightOccipitoCerebellar | only brainlife | right hemisphere occipito-cerebellar tract |
| **46** | rightParietoCerebellar | only brainlife | right hemisphere parieto-cerebellar tract |
| **47** | rightSLF1And2 | SLF_I_right\|SLF_II_right | right hemisphere superior longitudinal fasciculus I and II (combined) |
| **48** | rightSLF3 | SLF_III_right | right hemisphere superior longitudinal fasciculus III |
| **49** | rightTPC | only brainlife | right hemisphere temporo-parietal connection |
| **50** | rightThalamicoCerebellar | only brainlife | right hemisphere thalamo-cerebellar tract |
| **51** | rightUncinate | UF_right | right hemisphere uncinate fasciculus |
| **52** | rightVOF | only brainlife | right hemisphere vertical occipital fasciculus |
| **53** | rightbaum | only brainlife | right hemisphere Baum's loop (dorsal optic radiation) |
| **54** | rightcingulum | CG_right | right hemisphere cingulum bundle |
| **55** | rightfrontoThalamic | only brainlife | right hemisphere fronto-thalamic tract |
| **56** | rightmeyer | only brainlife | right hemisphere Meyer's loop (ventral optic radiation) |
| **57** | rightmotorThalamic | only brainlife | right hemisphere motor-thalamic tract |
| **58** | rightpArc | only brainlife | right hemisphere posterior arcuate fasciculus |
| **59** | rightparietoThalamic | only brainlife | right hemisphere parieto-thalamic tract |
| **60** | rightspinoThalamic | only brainlife | right hemisphere spinothalamic tract |
| **61** | righttemporoThalamic | only brainlife | right hemisphere temporo-thalamic tract |

The Brainlife bundle set extends beyond the TractSeg parcellation [2] by including several recently characterized white matter tracts not represented in TractSeg, most notably a comprehensive set of cerebellar tracts (motorCerebellar, parietoCerebellar, occipitoCerebellar, thalamicoCerebellar, anterioFrontoCerebellar, and their contralateral counterparts), subdivisions of the optic radiation (Meyer's loop and Baum's loop), finer subdivisions of the middle longitudinal fasciculus (MDLFang and MDLFspl), and posterior association tracts linking the dorsal and ventral visual streams (pArc, VOF, TPC). Furthermore, even for tracts that nominally exist in both frameworks, the two schemes do not define anatomically identical bundles, as they differ in their underlying segmentation philosophy: Brainlife WMA delineates bundles based on their full anatomical trajectory, including cortical projection zones, whereas TractSeg parcellates tracts based on the geometric position of midline crossing fibers or canonical ROI-based criteria. The corpus callosum is an example of this discrepancy: the anteriorFrontalCC bundle in Brainlife WMA most directly corresponds to TractSeg CC_1 (Rostrum) and CC_2 (Genu), both capturing interhemispheric prefrontal connections at the midline, while the forcepsMinor represents a broader spatial extent by additionally encompassing the fan-shaped projections radiating into the anterior frontal white matter and thus should not be directly equated with CC_1 and CC_2 alone. Similarly, the middleFrontalCC corresponds approximately to CC_3 (Rostral body), and CC_4 (Anterior midbody), and the forcepsMajor extends beyond CC_7 (Splenium) to include occipital projection fibers. The parietalCC bundle corresponds most closely to the anterior portion of CC_6 (Isthmus), capturing interhemispheric parietal connections while excluding the posterior auditory fibers also encompassed by CC_6, reflecting the fact that CC_6 straddles both parietal and auditory cortical territories, whereas parietalCC is restricted to the former.
